# Low-intensity vibration (LIV) cellular mechanotherapy reprograms tumor transcriptomes to suppress cancer-promoting inflammation

**DOI:** 10.64898/2026.04.07.716739

**Authors:** Stephene S. Meena, Benson K. Kosgei, Geofrey F. Soko, Wen Zheng, Shujiao Yu, Chuxiang Hu, Yuting Zhong, Binglin Li, Li Wang, Julius Mwaiselage, Xinju Hou, Ray P.S. Han

## Abstract

LIV is a non-pharmacological intervention for multiple pathologies, most notably pain, yet its potential in oncology remains unexplored due to poorly understood mechanotransduction pathways. Its therapeutic efficacy depends on precise dosing parameters - frequency, amplitude, and duration; suboptimal application, whether excessive or insufficient, risks exacerbating symptoms or rendering treatment ineffective. In a breast cancer–immune cell co-culture model, a 15-min regimen of 90-Hz, 0.43-g LIV attenuates protumorigenic crosstalk by suppressing proinflammatory cytokines and reprogramming malignant cells into a transient metastasis-incompetent “benignant” state that retains oncogenic markers but loses invasive capacity. Our results reveal LIV as a non-drug inhibitor of IFNγ and CXCL12β - silencing IFNγ impairs inflammatory bursts, while silencing CXCL12β starves the CXCR4/CXCR7 axis, depriving tumor cells of motility cues. LIV enforces a genome-wide transcriptional brake, upregulating tumor-suppressive miRNAs (hsa-miR-141, hsa-miR-122) while downregulating protumorigenic miRNAs (hsa-miR-105-5p, hsa-miR-181a-5p). This work repositions LIV as a non-drug modality for coordinately targeting multiple cancer hallmarks.

**Graphical abstract:** 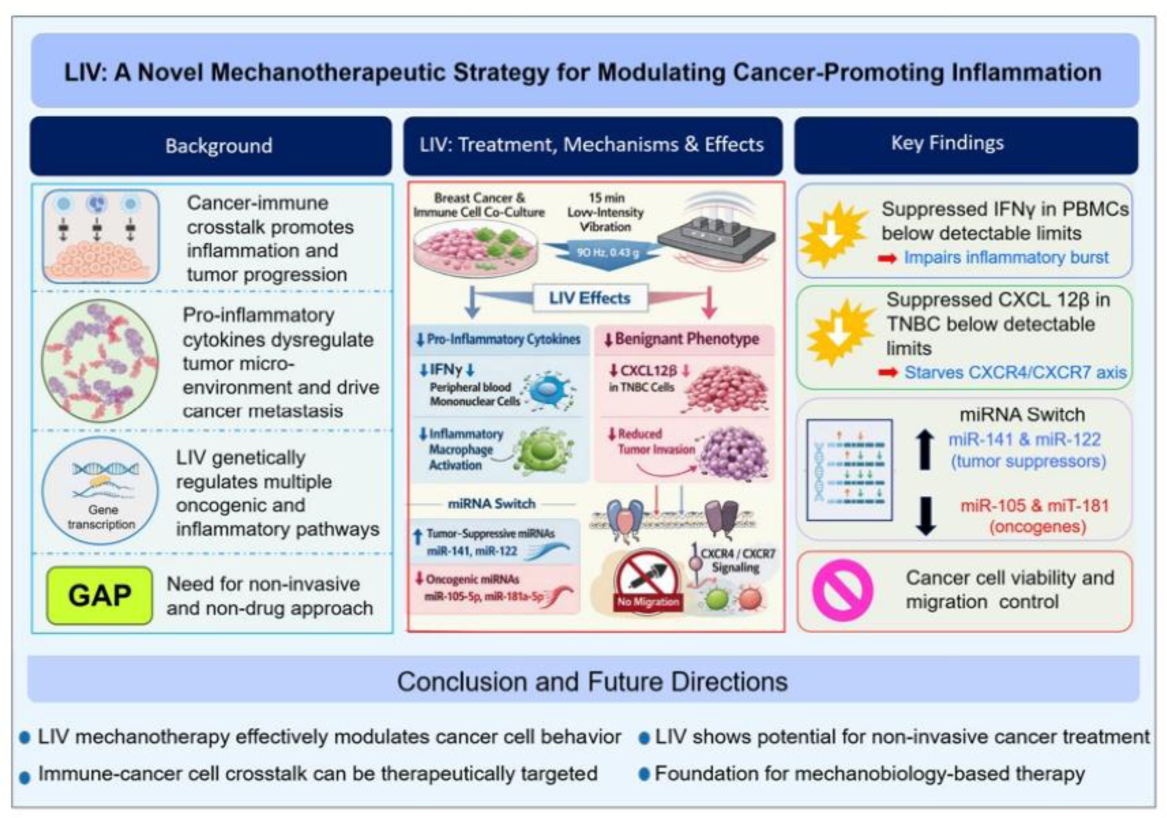

**Highlights:** *Dosing Precision:* LIV mechanotherapy has an optimal therapeutic window. Too little is ineffective, while too much can exacerbate symptoms. This mirrors the pharmacokinetic language of classical pharmacology.

*The “Benignant” Concept:* A 15-min regimen of 90-Hz, 0.43-g LIV attenuates protumorigenic crosstalk by suppressing proinflammatory cytokines and reprogramming malignant cells into a transient, metastasis-incompetent benignant state defined by mechanical constraint rather than pharmacological intervention.

*Dual Cytokine Silencing:* LIV acts as a non-pharmaco-logical inhibitor of IFNγ and CXCL12β. IFNγ is a double-edged sword in cancer (pro-immunity vs. exhaustion), and CXCL12β is the specific isoform implicated in metastasis. Targeting both simultaneously addresses immune crosstalk and chemotaxis.

**In brief:** Meena et al. demonstrate that the efficacy of LIV hinges on optimal dosing parameters. When applied correctly, it functions as a non-drug modality to target multiple oncogenic and inflammatory pathways. At the molecular level, LIV drives global transcriptional repression via miRNA regulation, increasing tumor-suppressive miRNAs (hsa-miR-141, hsa-miR-122) and decreasing protumorigenic miRNAs (hsa-miR-105-5p, hsa-miR-181a-5p).

## INTRODUCTION

Our expanding ability to decode cellular communication is fueling a new era of disruptive biomedical breakthroughs. At the heart of this revolution lies the intricate web of signaling networks that govern cellular behavior. Cells act as sophisticated sensors, using specialized membranes to detect and integrate a constant stream of multimodal cues—from chemical and mechanical to electrical and thermal signals. Remarkably, the cellular transduction apparatus operates with extraordinary fidelity in converting diverse extracellular inputs into intracellular biochemical events for processing in the nucleus. The output is then packaged into membrane-bound vesicles containing protein-ligand complexes to function as highly specific delivery vehicles, orchestrating intercellular dialogue with spatiotemporal accuracy. By mastering this fundamental code of life, we are now engineering revolutionary therapies, ranging from exosome-based delivery systems to sophisticated synthetic signaling circuits, to intervene in disease at its most basic level.

Cellular signal detection is mediated by sensitive transmembrane protein receptors that initiate intracellular signal transduction cascades, ultimately altering cell behavior.^1–3^ This conversion of extracellular stimuli and eliciting defined biochemical responses is fundamental to cell communication. For example, electrical signals are converted by voltage-gated channels,^4^ while mechanical cues are transduced by stretch-activated ion channels^5,6^ to trigger conformational changes in mechanosensitive proteins.^7–10^ Beyond rapid signal relay, sustained mechanical loading can induce cytoskeletal contraction, directly transmitting physical forces to the nucleus. There, they remodel three-dimensional chromatin architecture and modulate the accessibility of DNA regulatory elements, thereby linking mechanical cues to transcriptional programs.^11^ Given this profound influence on cellular physiology, mechanical signaling is now recognized as a pivotal regulator in both tissue homeostasis and disease pathogenesis.^12^ Indeed, all forms of cellular signaling can modulate cell behavior and responses, offering a rich landscape of potential therapeutic targets. However, current therapies remains predominantly reliant on chemical interventions through drugs and biologics^13^ and overlooking the therapeutic potential of physical cues. Elucidating how cells perceive, integrate, and respond to multimodal signals is therefore essential for deciphering the pathophysiological mechanisms that drive disease progression. This paradigm is especially critical for understanding and treating diseases driven by chronic inflammation, including cancer.^14^

Inflammation, while a critical defense mechanism, is hijacked by cancer cells to promote survival, proliferation, and metastasis.^15,16^ This cancer-promoting inflammation (CPI) reprograms immune cells like macrophages into tumor-associated macrophages (TAMs), which can constitute up to 50% of a solid tumor mass.^17,18^ TAMs create a pro-tumor environment by driving angiogenesis, suppressing anti-tumor T-cells, and secreting pro-inflammatory cytokines that exacerbate the disease.^17,19–23^ Despite this well-established role of inflammation in tumor initiation, progression, invasion, recurrence, metastasis, and resistance to treatment,^16,20^ current anticancer therapies largely overlook CPI management. Further, pharmacological interventions are often inadequate because their single-target approach cannot effectively disrupt the complex, interconnected signaling network that orchestrates CPI. Emerging evidence suggests that mechanical cues offer a promising alternative by simultaneously modulating multiple pathways.^21,22^ Macrophages, as first responders, are profoundly mechanosensitive, dynamically altering their functional polarization in response to external physical cues via integrin-mediated mechanotransduction.^23–26^ For example, low-intensity vibration (LIV) has been shown to suppress inflammatory activity by downregulating proinflammatory cytokines like IL-1β and TNFα while upregulating the anti-inflammatory cytokine IL-10.^27^ However, while the anti-inflammatory potential of mechanical cues is clear, their role within the tumor-immune microenvironment (TIME) remains poorly understood. Specifically, how mechanical modulation influences CPI activity is a critical unanswered question.

This study investigates how mechanical stimulation disrupts cancer progression by reprogramming cellular communication within TIME. Using a breast cancer-immune cell co-culture model, we tested the hypothesis that a 15-min, 90-Hz, 0.43-g LIV protocol exerts its effect through two synergistic mechanisms: suppressing pro-inflammatory cytokine (PIC) secretion via immune cell modulation, and inducing a phenotypic transformation of malignant cells into a metastatically incompetent state. We term this transformed state *benignant*—a benign-like condition where cells retain transformation markers but lose metastatic capacity. Our data supports this hypothesis, showing that LIV stimulation of breast cancer cells (MDA-MB-231) cells co - cultured with peripheral blood mononuclear cells (PBMCs) significantly downregulates key immunomodulators, reduces pro-inflammatory mediators (PICs, chemokines, and oncogenic microRNAs), and decreases cancer cell viability. These findings position LIV as a promising mechanotherapeutic intervention capable of simultane ously attenuating pro-tumorigenic signaling and curbing cancer aggressiveness.

## RESULTS WITH BRIEF COMMENTARIES

### Development and calibration of a tunable mechanotransduction system

To investigate LIV effects in cancer and immune cells, we developed a tunable vibration platform that delivers precisely calibrated mechanical stimuli to cell cultures (Figure 1A). The system was validated using MDA-MB-231 cells and PBMCs (Figure 1B). Following calibration, the actuator transmitted discrete frequencies (90, 150, and 300 Hz) set by interchangeable gears with an inverse frequency–displacement relationship alongside proportional increases in acceleration and velocity. At 90 Hz frequency, measured acceleration, velocity and displacement amplitude were: 4175 mm/s² (0.43 g), 2.825 mm/s, and 0.03 mm, respectively (Figures 1C, S1 and Table S1). These mechanical signatures setup the biophysical boundary conditions for LIV-based cellular mechanostimulation. The platform offers a reproducible test -bed to decode how me chanical cues curbs cancer progression and prototype novel mechano transduction-targeted therapies.

**Figure 1.**
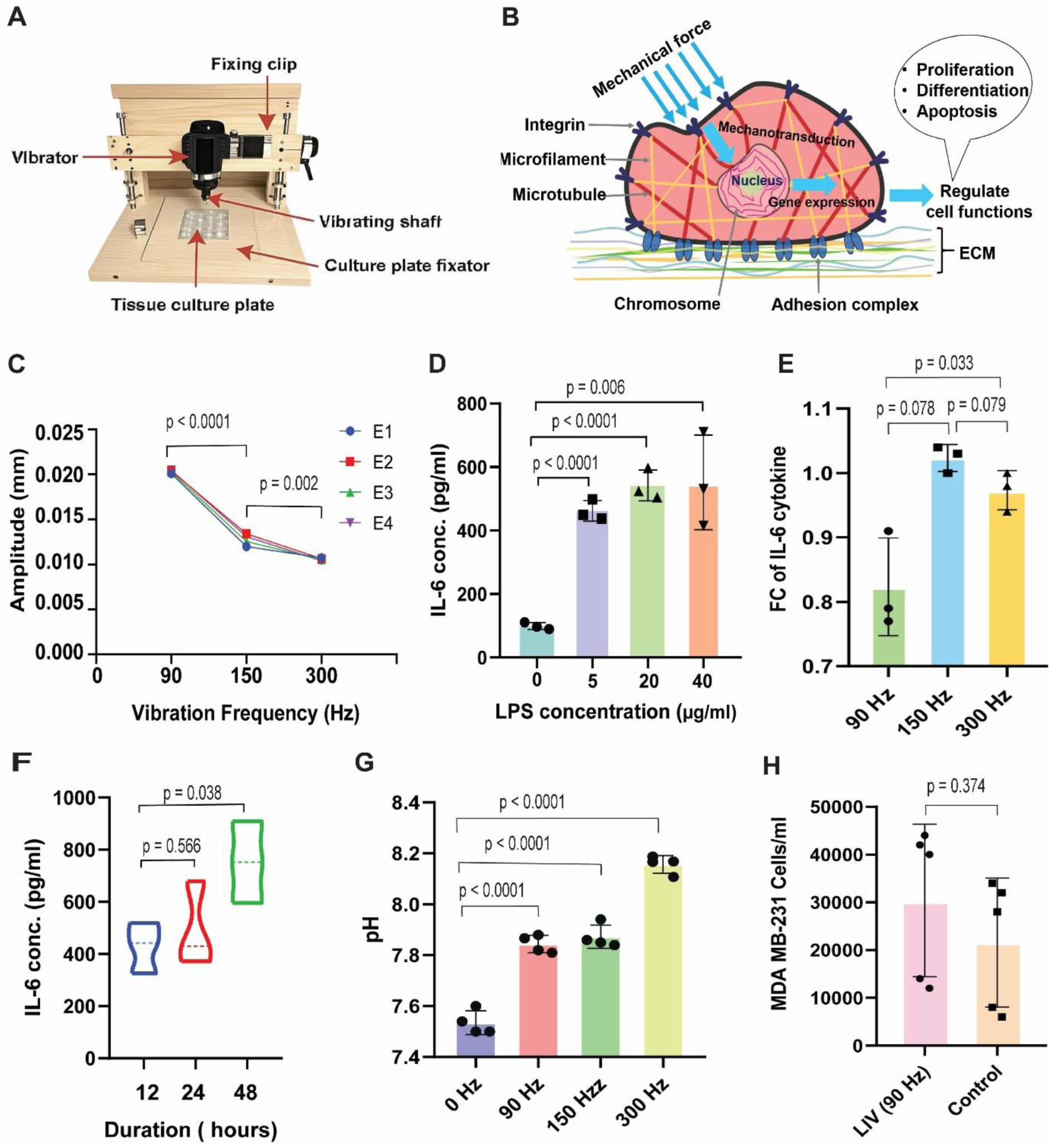
System calibration, mechanotransduction mechanism, and cellular effects of LIV. (A) Schematic of the custom-built vibration system, which uses a rigidly mounted gun with adjustable stoppers for tuning vibration amplitude to ensure reproducible mechanical stimulation. (B) Overview of mechanotransduction: extrinsic mechanical forces are converted into biochemical signals that activate downstream pathways, leading to changes in gene expression, protein synthesis, and cell phenotype—thereby, regulating cellular proliferation, differentiation, and apoptosis. (C) Calibration of vibration frequencies at different gear settings: gears 1, 2, and 4 corresponded to 90 Hz, 150 Hz, and 300 Hz, respectively, measured at the plate bottom. Vibration frequency varied inversely with displacement amplitude (*n* = 4; *p* < 0.05). (D) Supernatant analysis of LPS-stimulated PBMCs showed a linear relationship between LPS concentration (0– 10 µg/ml) and IL-6 secretion (*p* < 0.05). Maximal PBMC proliferation was observed at 5 μg/ml of LPS (*n* = 3). (E) Fold change (FC) of IL-6 cytokine secreted by vibrated PBMCs showed most reduction at the 90 Hz LIV (*n* = 3). (F) Time-course of IL-6 secretion from LPS-stimulated PBMCs revealed peak cytokine levels at 48 h (*n* = 3). (G) LIV increases the supernatant pH of LPS-stimulated PBMCs in a frequency-dependent manner (*n* = 4). (H) LIV did not significantly alter the proliferation of MDA-MB-231 cells compared to static controls (*n* = 5; *p* = 0.374).

### Baseline inflammatory response in LPS-activated PBMCs under static conditions

To evaluate how mechanical vibration modulates IL-6 release, we first mapped the baseline inflammatory response of bacterial lipopolysaccharide (LPS)-activated PBMCs under static conditions. The objective of this experiment was to establish the optimal LPS concentration for stimulating PBMCs. The experiments revealed a classic dose-response curve. A robust, approxi mately linear relationship (*R*²= 0.92) between LPS dose and IL-6 secretion was observed within mid-range concentrations of 5-40 μg/ml (Figure 1D). However, this linear trend was constrained by threshold effects. At low concentrations (<5 μg/ml), LPS was insufficient to trigger a significant inflammatory response. Conversely, at high concentrations (> 40 μg/ml), the system became saturated, resulting in a plateau and subsequent decrease in IL-6 production.

### Our LIV protocol

To identify a vibration frequency that effectively suppresses PIC secretion, we evaluated the fold change (FC), which is defined as the ratio of IL-6 levels in vibrated to non-vibrated PBMCs across three frequencies: 90 Hz, 150 Hz, and 300 Hz. IL-6 secretion showed the most reduction at 90 Hz (Figure 1E), establishing this frequency as optimal. Summarizing, our LIV protocol is defined by 15-min, 90 Hz and 0.43 g.

### Temporal dynamics of LPS-induced IL-6 release

To define the optimal time window for intervention, we performed a temporal analysis of IL-6 secretion kinetics in LPS-stimulated PBMCs. Cell culture supernatant of activated PBMCs were sampled at 12, 24 and 48 h. IL-6 rose steadily from 428.87± 97.51 pg/mL (12 h) to 493.89 ± 163.41 pg/mL (24 h) and peaked at 752.50± 156.58 pg/mL (48 h) post-stimulation (Figure 1F). This sustained upswing defines the 24–48 h window as the optimal intervention period for timing mechanomodulatory interventions. These findings complement our frequency-response data by establishing both the intensity and temporal parameters of LPS-induced inflammation in PBMCs, providing a complete set of data for systematically investigating how mechanical cues can reprogram immune cell activity.

### Impact of LIV on pH and cancer cell growth

Our data showed that LIV generated a frequency-dependent alkalinization of conditioned medium from LPS-activated PBMCs, with the largest pH rise at 150–300 Hz (Figure 1G). The shift may reflect vibration-induced changes in cellular metabolism and/or altered secretion of acidic metabolites, potentially influencing the behavior of both cancer and immune cells by modulating their extracellular milieu.

When MDA-MB-231 cells were intermittently exposed to 90 Hz LIV over a 24-h duration, they showed a mild proliferative response to mechanical stimulation compared to static controls (*p* = 0.374; Figure 1H), a finding consistent with published reports.^28,29^ These results highlight both direct and indirect effects of LIV, setting the stage for more complex experiments.

### LIV modulates migration, proliferation, and viability in tumor-immune co-cultures

The dysregulation of core cellular processes—migration, proliferation, and viability—is a hallmark of cancer. Here, we demonstrate that LIV significantly modulates these processes in a model of tumor-immune interactions using co-cultures of MDA-MB-231 cells and PBMCs. We employed both 2D and 3D platforms, including a Transwell system to isolate and capture paracrine signaling effects. We captured fluorescence images of MDA-MB-231 cells and PBMCs (Figures 2A and 2B), and fluorescence and bright-field images of spheroids generated by MDA-MB-231 cells and PBMCs (Figures 2C and 2D). To interrogate cell motility a confluent 2-D tumor–immune co-culture of MDA-MB-231 cells and PBMCs (Figure 2E) was scratch-wounded with closure monitored over time (Figures S2, S3 and S4). Our results reveal that LIV exerts potent anti-tumor effects.

**Figure 2.**
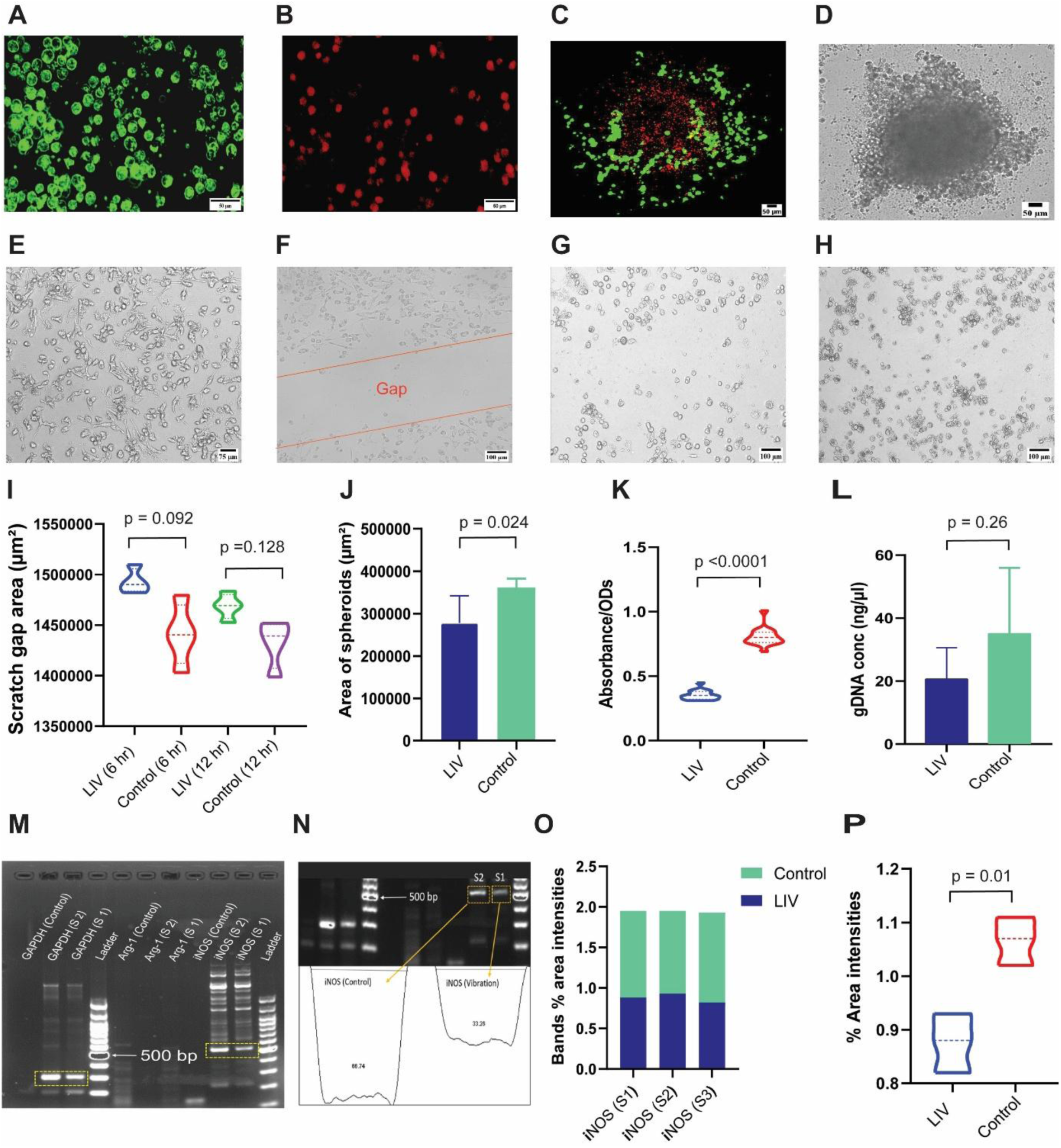
LIV suppresses cancer cell viability and migration, and restricts cell growth in 2D and 3D co-culture models, respectively. (A-B) Representative fluorescence images showing (a) MDA-MB-231 cells stained with green CellTracker and (b) PBMCs stained with red CellTracker. (C) Fluorescence image of a co-culture spheroid of MDA-MB-231 cells (green) and PBMCs (red). (D) Bright-field image of a co-culture spheroid of MDA-MB-231 cells and PBMCs. (E) Bright-field image of a monolayer co-culture of MDA-MB-231 cells and PBMCs. F-H) Scratch wound images of the co-culture taken at (f) 0h, (g) 6h, and (h) 12h. (I) Wound-closure assay quantitation. Control cells exhibited faster migration than mechanically stimulated (90 Hz) cells, as shown by scratch-closure area over time (*n*=4). (J) Mean spheroid size from co-cultured MDA-MB-231 cells and PBMCs was reduced under 90Hz vibration compared with the static control (0Hz) (*n*=5; *p*<0.05). It is obvious that mechanical stimulation suppressed spheroid growth. (K) Cell viability of MDA-MB-231 cells was significantly lower in the 90Hz group (43.8±5.1 %) than in the control (L) Genomic DNA content of PBMCs was lower in vibrated samples than in controls (*n*=4). (M) Separation of PCR amplicons by agarose gel electrophoresis. (N) Densitometric analysis (ImageJ) of agarose gel bands. Normalized percentage area intensities are shown for iNOS amplicons in vibration vs. control groups (*n*=3). (O) Normalized percentage comparisons for independent experiments (*n*=3). (P) Boxplot of normalized mean percentage area intensities (*p*<0.05).

#### Migration

In scratch-wounded 2-D co-cultures, LIV delayed wound closure; quantitative ImageJ tracking showed a consistent (non-significant) reduction in cancer-cell migration at 6 h and 12 h (Figures 2F, 2G and 2H). Quantitative analysis confirmed a consistent suppression of MDA-MB-231 migration in the presence of PBMCs (Figure 2I), indicating a trend toward LIV-induced inhibition of motility. *3-D growth*: LIV markedly restricted spheroid expansion of MDA-MB-231 cells co-cultured with PBMCs (*p* < 0.05; Figure 2J), indicating potent anti-proliferative activity in a physiologic 3-D milieu. *Viability*: LIV reduced 2-D monoculture viability to 43.8± 5.1 % versus controls (*p* < 0.0001; Figure 2K) and triggered pronounced total gDNA loss (Figure 2L), confirming a genomic modulation of PBMCs.

Concluding, LIV acts as a multi-faceted modulator of tumorigenic processes, impairing migration, proliferation, and viability in both direct and indirect co-culture of breast cancer cells with immune cells.

### LIV suppresses inflammatory response

To assess the anti-inflammatory potential of LIV, we investigated its capacity to modulate macrophage polarization within cultured PBMCs. We quantified the expression of inducible nitric oxide synthase (iNOS) and arginase-1 (Arg-1), canonical markers of pro-inflammatory (M1) and anti-inflammatory (M2) phenotypes, respectively.^30^ In Transwell experiments, pairing MDA-MB-231 cells with PBMCs, LIV provoked a pronounced shift away from the pro-inflammatory M1 phenotype: gDNA analysis revealed a marked suppression of iNOS expression in vibrated PBMCs (Figures 2M, 2N, 2O and 2P). In contrast, Arg-1 remained undetectable in both vibration and static co-cultures (Figures 2M, 2N, 2O and 2P). These data indicate that LIV selectively attenuates M1 polarization without driving overt M2 commitment, effectively dampening the inflammatory milieu. Primer sequences for iNOS, Arg-1, and the housekeeping control gene (GAPDH) are listed in Table S2.

### LIV reprograms inflammatory secretomes

Building on the temporal dynamics of PIC secretion by LPS-stimulated PBMCs, we asked whether LIV could reprogram the secretome output from both immune and cancer cells. Supernatants from 90-Hz-vibrated monocultures and co-cultures were profiled by ELISA. The results are as follows. Immune cells: LIV significantly altered the secretory profile of LPS-activated PBMCs, suppressing key PICs−IL-6 and IFNγ (Figures 3A and 3B) while promoting the release of the anti-inflammatory cytokine IL-10 and the monocyte-recruiting chemokine MCP-1 (Figure 3C and 3D). Cancer cells: a parallel effect was observed in MDA-MB-231 breast cancer cells, where LIV treatment notably inhibited the secretion of pro-inflammatory factors, includ ing IL-6, TNFα, and the metastasis-promoting chemokine CXCL12β (Figures 3E, 3F and 3G).

**Figure 3.**
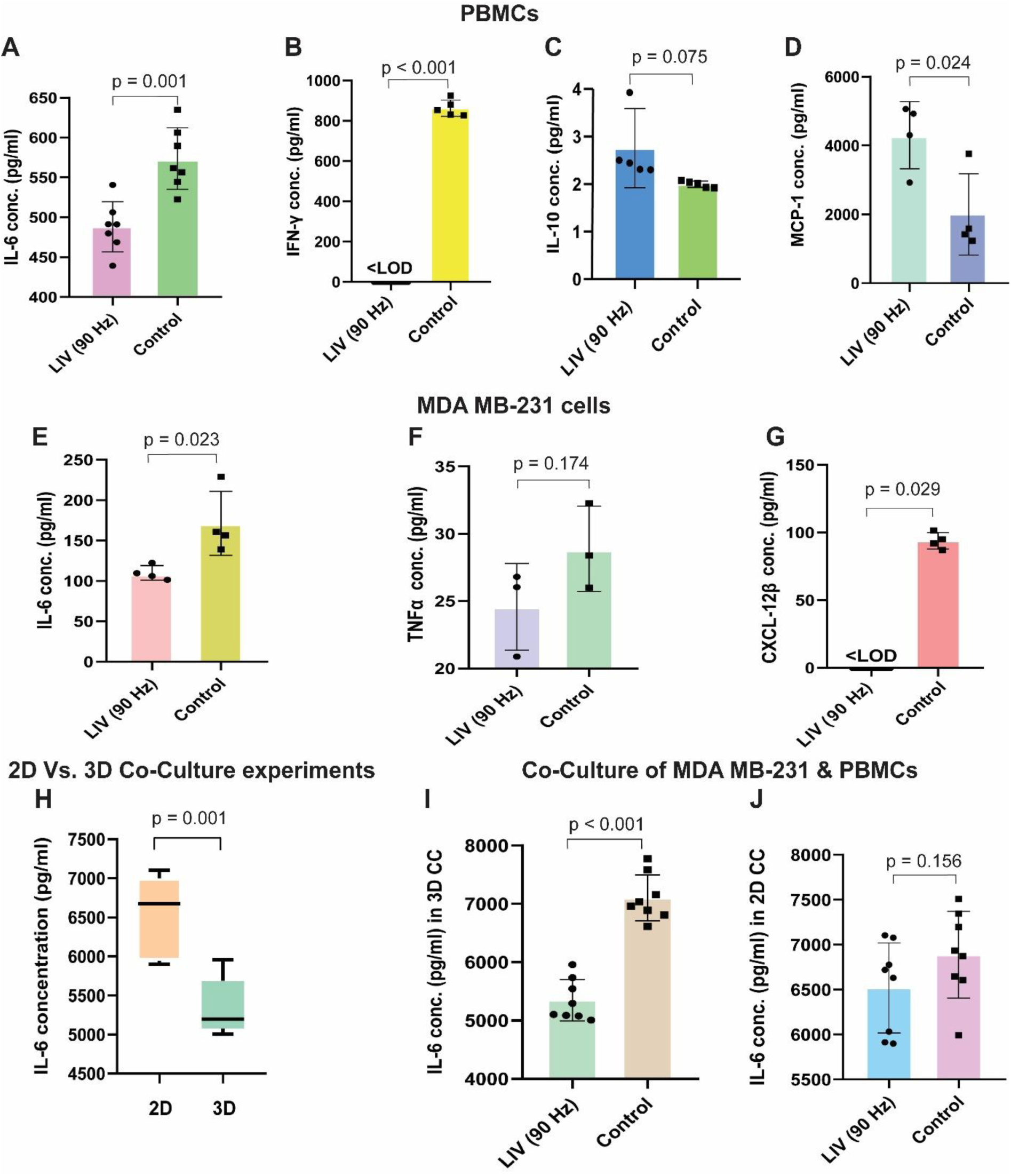
LIV modulates cytokine secretion in immune and cancer cells. Cytokine concentrations in cell culture supernatants were quantified by ELISA (mean ± SD) and statistical significance was determined by Student’s t-test. (A) IL-6 concentration in vibrated PBMCs was significantly reduced compared to static control (488.26±31.49 pg/ml vs. 573.76± 38.66pg/ml, *p* = 0.001, *n* = 7). (B) IFNγ in LIV-treated cells was below the lower limit of detection (LOD) compared to static control concentration of 854.41± 36.01 pg/ml, *p* < 0.001, *n* = 5). (C) IL-10 concentration in vibrated PBMCs showed a non-significant increase over control (2.8± 0.83 pg/ml vs. 2.0± 0.70 pg/ml, + 40 %; *p* = 0.08, *n* = 5). (D) MCP-1 concentration increased markedly in vibrated PBMCs (4305± 975 pg/ml vs. 2001± 1183 pg/ml, + 115 %; *p* = 0.02, *n* = 4). (E) IL-6 in vibrated MDA-MB-231 cells was significantly lower than in controls (109.93± 9.03 pg/ml vs. 171.35± 39.61 pg/ml, − 36 %, *p* = 0.02, *n* = 4). (F) TNF-α levels did not differ significantly between vibrated and control MDA-MB-231 cells (24.57± 3.22 pg/ml vs. 28.88± 3.17 pg/ml, − 15 %; *p* = 0.17, *n* = 3). (G) CXCL-12β in LIV-treated MDA-MB-231 cells was below the LOD, whereas controls measured at 56.25 ± 39.40 pg/ml (*p* = 0.03, *n* = 4). (H) In co-culture models, IL-6 levels were higher in 2D than in 3D spheroids under LIV (6518± 501 pg/ml vs. 5351± 355 pg/ml, − 22%; *p* = 0.001, *n* = 8). (I) IL-6 concentration in vibrated 3D co-cultures was significantly reduced compared with static controls (5351± 355 pg/ml vs. 7104± 392 pg/ml, − 25 %, *p* < 0.001, *n* = 8). (J) IL-6 levels in 2D co-cultures did not differ significantly between vibrated and control groups (6518± 501 pg/ml vs. 6887± 483 pg/ml, −5%; *p* = 0.16, *n* = 8).

LIV-mediated inflammatory suppression is even more remarkable in co-cultured MDA-MB-231 and PBMC 3D spheroid experiments (Figure 3H). In the 3D co-culture, 90-Hz LIV suppresses IL-6 secretion by > 25 % (p< 0.001; Figure 3I), whereas the same treatment in 2D co-culture yielded only a non-significant downward trend (p= 0.156; Figure 3J). Full statistical details are provided in Tables S4 (cytokine assays) and S5 (LPS titration controls).

### LIV modulates EV biogenesis and molecular cargo of MDA-MB-231 cells

LIV induced significant changes in both physical properties and molecular cargo of MDA-MB-231-derived extracellular vesicles (EVs). After confirming their exosomal identity through size, morphology, and biochemical marker expression (Figures 4A, 4B and 4C), we investigated how LIV affected their production. Nanoparticle-tracking analysis (Figure 4D) revealed that LIV treatment profoundly altered EV biogenesis. LIV mechanically stimulated cells to secrete large quantities of EVs (Figure 4E), LIV almost tripled the yield of small EVs compared to static control (11. 2± 0.37× 10⁹ particles/mL vs. 4.0 3± 0.2 3× 10⁹ particles/mL; *p* < 0.001 with a dramatic shift towards smaller particles (137.5± 1.5 nm vs. 18 2± 6.4 nm;*p* < 0.001) (Figure 4F). Zeta potential analysis showed that the s urface charge of the two groups stayed same (−12. 6± 1.1 mV vs −12. 7± 0.9 mV, *p* = 0.92, Figure 4G) despite a surge in small EVs. Intriguingly, this EV surge did not extend to miRNA cargo. LIV -treated cells actually secreted less tumor-associate miRNAs, both in free form and encapsulated within EVs than control (Figure 4H). Across all samples, miRNAs were predominantly vesicle-bound rather than freely -distributed in the extracellular milieu. On average, the total con centration of encapsulated -miRNAs in tumor-derived exosomes was 3.9-fold higher than free -miRNAs in the culture medium (36.72± 1.78 ng/µl vs 9.37± 1.20 ng/µl, *p* = 0.0002, Figure 4H).

**Figure 4.**
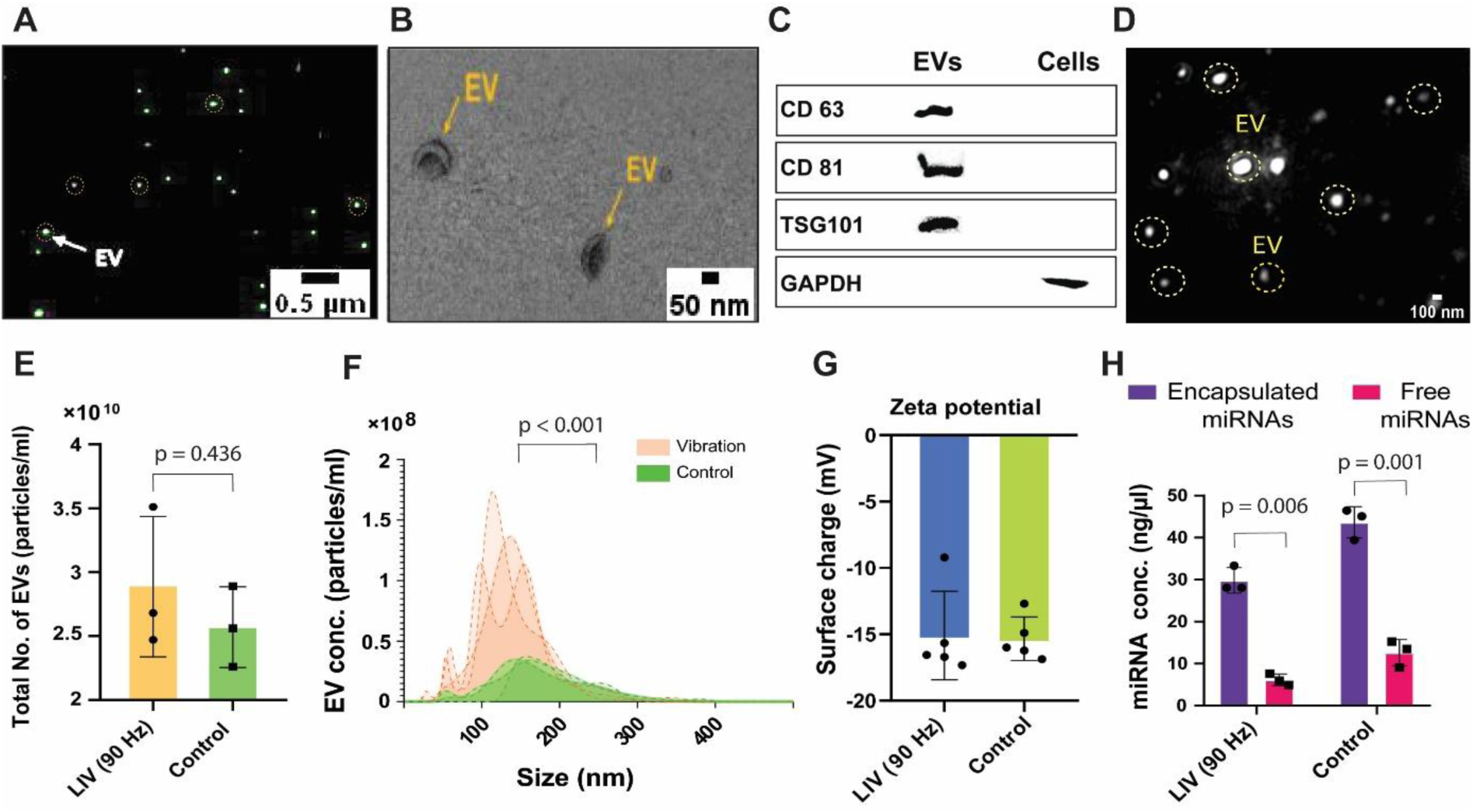
Characterization of EVs. (A) Confocal fluorescence image of PKH67-stained EVs (green). (B) Transmission electron micrograph (TEM) of MDA-MB-231-derived EVs. (C) Western blot confirming EV identity via the detection of tetraspanins CD63, CD81, and TSG101, with GAPDH as a loading control. (D) Representative nanoparticle tracking analysis (NTA) frame view showing EV size distribution and integrity. (E) Total EV concentration secreted by vibrated MDA-MB-231 cells versus static control (2.89± 0.32 × 10^10^ vs. 2.50 ± 0.18 × 10^10^ particles/ml, *p* = 0.436, *n* = 3). (F) NTA comparison of tumor-derived EVs (TDEs): LIV-treated cells secreted smaller EVs (137.5± 1.5 nm vs. 182± 6.4 nm; concentration at mean size: 11.2 ± 0.37 × 10⁹ particles/mL vs. 4.03 ± 0.23 × 10⁹ particles/mL; *n* = 3). (G) Zeta-potential analysis showed no significant difference in the surface charge of EVs released by vibrated MDA-MB-231 cells vs static cells (*p* = 0.92, *n* = 3). (H) Comparison of miRNA content in TDEs versus free miRNAs in cell-culture medium (CCM). Encapsulated miRNA concentration in TDEs was 3.91-fold higher than free miRNA in CCM (36.72± 1.78 ng/µl vs. 9.37± 1.20 ng/µl, *p* = 0.0002, *n* = 3). Both encapsulated and free miRNA levels were lower in LIV-treated cells than in controls (*n* = 3).

### LIV drives a genome-wide suppressive program in MDA-MB-231 cells

Transcriptomic profiling of triple-negative breast cancer cells was performed to decipher the molecular basis of LIV’s effects. RNA-seq revealed the dominant signature was a genome-wide suppression of the transcriptome in LIV-treated cells, characterized by a significant reduction in upregulated protein-coding genes relative to static controls (Figure 5A). Small-RNA libraries were dominated by rRNA and tRNA species for both groups (Figure 5B). High-confidence miRNA–mRNA interactions were extracted by intersecting TargetScan, miRanda and RNAhybrid predictions, yielding 3.8 million consensus targets (Figure 5C). Pearson correlation analysis heat-maps confirmed that LIV induces a distinct transcriptional state (Figure 5D). Replicates within the control (C1-C3; *r* = 0.9-1.0) and LIV-treated (V1-V3; *r* = 0.7-1.0) groups showed strong intra-group clustering, validating experimental reproducibility and the significant impact of mechanical stimulus. A core set of 265 genes remained unchanged across controls (Extended Data Figure5). Although only 13 of 717 detected miRNAs were differentially expressed (seven up, six down; Figures 5E and 5F), genome-wide DESeq2 profiling demonstrated broad transcriptional suppression, with LIV replicates (V1-V3) displaying uniformly lower expression than controls (C1-C3; Figure 5G). Subsequent qPCR validation on EVs derived from these cells (custom TaqMan primers, Table S6; ΔΔCt calculations, Table S7) validated these changes: LIV significantly downregulated onco-miRs hsa-miR-105-5p and hsa-miR-181a-5p while upregulating tumor-suppressors hsa-miR-141 and hsa-miR-122 (*p* > 0.05, Figure 5H).

**Figure 5.**
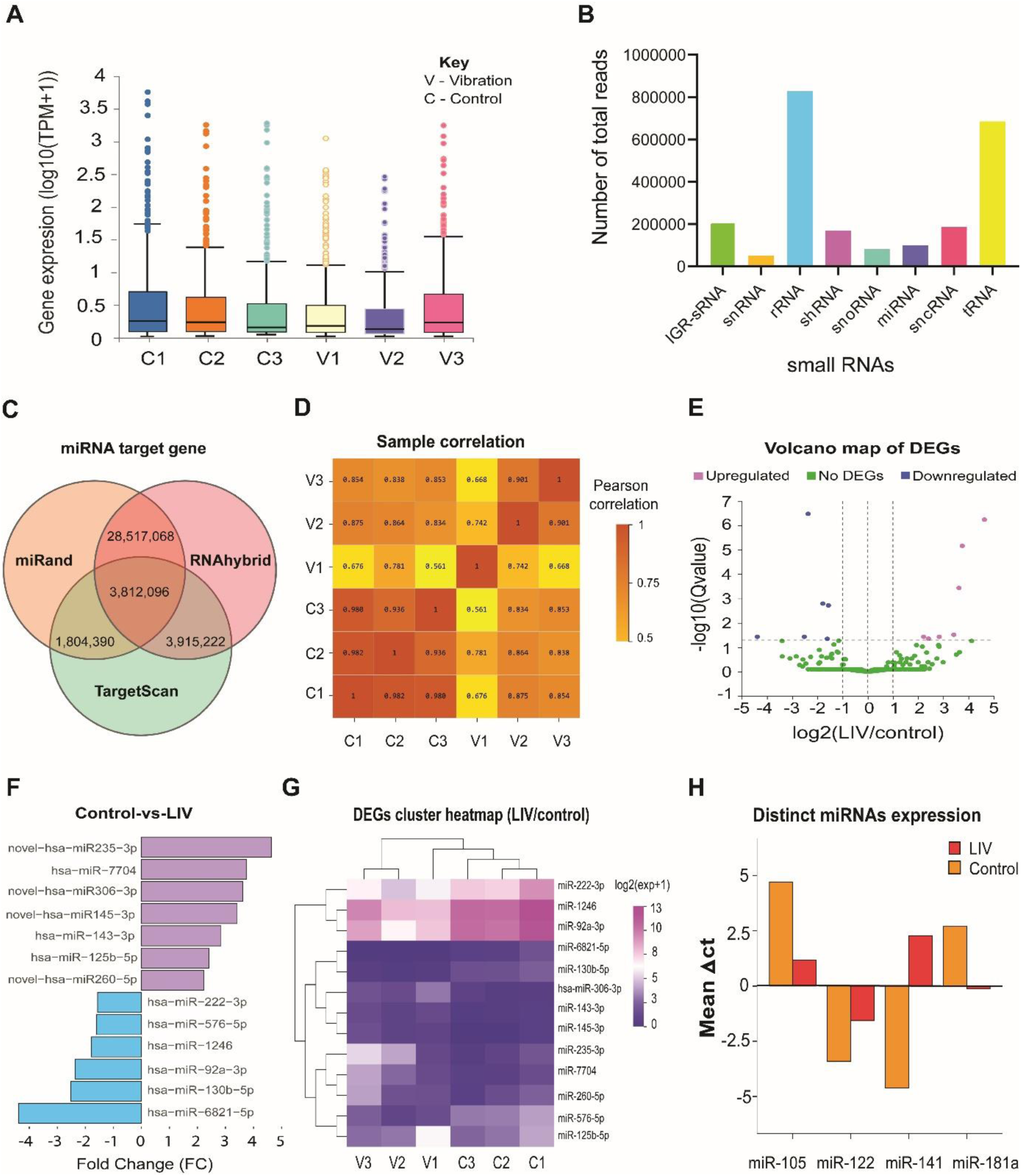
LIV reprograms the miRNA expression profile. (A) Box-plot of gene-expression distribution across sequenced samples (C1-C3: control; V1-V3: LIV-treated), revealing an overall higher expression in control vs. vibration groups. (B) Small RNA-seq profiling identified small hairpin RNA as the most abundant RNA class, followed by miRNA and other non-coding RNA species. (C) Venn diagram depicting the overlap of predicted gene targets for differentially expressed miRNAs in MDA-MB-231 cells using three independent algorithms: miRand, RNAhybrid, TargetScan (predicted targets, *p* < 0.05). Each circle represents the target relationship (miRNA-gene pairing relationship) predicted by an algorithm, and the overlapping area denotes the common target relationship of the three algorithms taken together. (D) Direct comparison of gene-expression levels between vibrated samples (V1–V3) and non-vibrated controls (C1–C3). (E) Volcano plot showing that 13 miRNAs were differentially expressed in LIV-treated samples out of 717 detected (DEseq2 criteria: [log2FC]> 0 & *Q* < 0.05). (F) Differentially expressed miRNAs in control vs. vibration are shown as up-regulated genes (purple) and down-regulated genes (blue) (Q < 0.05). (G) Heatmap of miRNA-sequencing data showing a cluster of DEGs with lower expression in vibrated samples (V1–V3) than in controls (C1–C3). Color gradient from purple (high) to blue (low) corresponds to high-to-low levels of expression. (H) RT-qPCR validation of exosomal miRNAs: EVs from LIV-treated MDA-MB-231 cells downregulated hsa-miR-105 and hsa-miR-181a, and upregulated tumor-suppressors hsa-miR-141 and hsa-miR-122 (*p* > 0.05).

Gene abundance trends were visualized with an RNA density map (Figure S6), and the chromosomal locations of genes targeted by differentially expressed microRNAs were mapped (Figure S7). Notably, our analysis revealed that a single microRNA, such as the differentially expressed hsa-miR-222-3p in the vibrated group, can regulate multiple transcripts (Figure S8).

### Gene-expression architecture and functional wiring of mechanosensitive transcriptomes

Gene levels across all samples were binned into three expression stringencies: Transcripts Per Million, TPM≤ 1 (silent), 1–10 (weak) and ≥ 10 (active) (Figure 6A) and DEGs in our dataset (DESeq2, padj< 0.05, |log₂FC|> 1) were interrogated with GO/KEGG (Q< 0.05). Cellular-compartment GO terms centered on nucleus, cytoplasm, cytosol, nucleoplasm, cytoskeleton and cell projection (Rich Ratio 0.7–1.0, Figure 6B), indicating that LIV targets the physical and regulatory core of the cell. KEGG pathways converged on cancer hubs, focal adhesion, cytoskeletal regulation, breast cancer, miRNAs in cancer, and mechanotransduction cascades (Rap1, Hippo, Oxytocin, MAPK, Ras, mTOR, Wnt; RichRatio 0.8–1.0, Figure 6C). Partitioning GO into biological process (BP), cellular component (CC) and molecular function (MF) (top-10 by gene count; Figure 6D) showed BP dominance (cellular process, biological regulation, and metabolism), indicating a widespread transcriptional response to mechanical cues. CC followed with genes mapping to cell, cell-part and organelle, whereas MF displayed the smallest set enriched for binding and catalytic activity. KEGG super-categories (Figure 6E) placed “signal transduction” at the apex across all classes; within organismal systems–the immune, endocrine and nervous systems prevailed, whereas for human diseases–cancers, infectious diseases and neurodegeneration topped the list. Under environmental-information processing, the most annotated pathways were signal transduction, signaling molecules and interaction, and folding, sorting and degradation.

**Figure 6.**
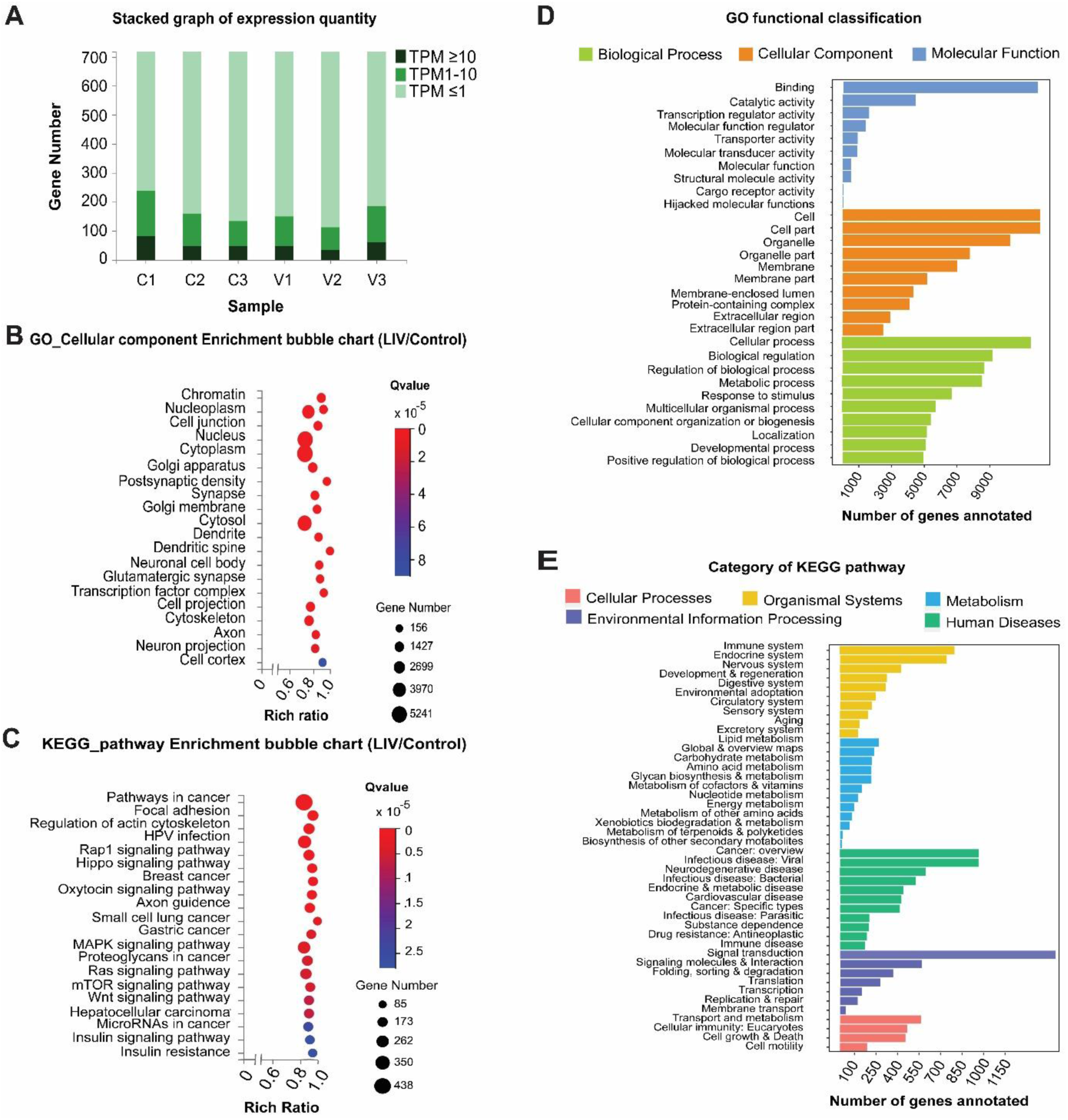
GO and KEGG enrichment and pathway analyses. (A) Density plot showing the distribution of gene expression levels (TPM) for each sample, confirming consistent data quality. Sample name and number of genes are listed in *x*- and *y*-axes, respectively, with color intensity representing expression level. (B) Top 20 significantly enriched Gene Ontology (GO) cellular-component terms among DEGs in vibration versus control (*Q* < 0.05). Bubble size corresponds to the number of genes per term; color denotes the *Q*-value (deep red: lower *Q*-value; blue: higher *Q*-value). (C) Top 20 enriched Kyoto Encyclopedia of Genes and Genomes (KEGG) pathways in vibration and control, and ranked by *Q*-value. Bubble size indicates the number of genes mapped to each pathway; color represents FDR-adjusted *p*-value (*Q*-value), designated with deep red a smaller *Q*-value, and blue a larger *Q*-value. (D) GO functional classification of DEGs between vibration and control groups. Bars represent the number of DEGs assigned to biological process (BP, blue), cellular component (CC, orange), and molecular function (MF, light blue) categories. (E) KEGG pathway classification using DEGs of vibration vs. control, summarized into six major categories: cellular processes, environmental information processing, genetic information processing, human diseases, metabolism, and organismal systems, showing the number of annotated genes in each.

## DISCUSSION

The recruitment of immune cells to sites of infection or injury is a cornerstone of inflammatory healing. However, tumors co-opt inflammatory processes to promote their growth and survival, in part by recruiting and modulating immune cells like macrophages. We demonstrated that the 15-min, 90-Hz LIV can fundamentally disrupt this protumorigenic crosstalk, leading to the suppression of tumor-promoting inflammation.

A key finding is LIV potently suppressed inflammatory response in LPS-activated PBMCs; in particular, reducing the secretion of a key cytokine interferon-γ (IFNγ) to undetectable levels, compared to 854.41± 36.01 pg/ml in controls (*p* < 0.001, *n* = 5, Figure 3B). The near-total abrogation suggests that LIV could act as a promising *non-drug inhibitor* of IFNγ and CXCL12 to curb pro-inflammatory signaling. This finding holds significant implications for macrophages, which are tissue-resident first responders for immune and inflammatory response, tissue repair, homeostasis, and tumor immunosurveillance.^31–34^ Macrophages display remarkable plasticity, polarizing along a spectrum of functional states in response to microenvironmental cues. While the classical inflammatory M1 state (induced by LPS/IFNγ) and the resolving stimuli M2 state (induced by, e.g. IL-4), represent two polarized extremes on a spectrum of macrophage responses,^35^ *in vivo* populations exhibit a complex continuum of activation states with profound consequences for disease and homeostasis.^36,37^ Our data suggest that LIV may fundamentally shift this polarization landscape.

LIV also silenced a key driver of metastasis within the cancer cells themselves. In MDA-MB-231 cells, LIV treatment significantly suppressed the chemokine CXCL12β, reducing its levels below LOD compared to 56.25± 39.40 pg/ml in controls (*p* = 0.03, *n=* 4, Figure 3G). Among the six known human splicing variants (CXCL12α-φ),^38,39^ CXCL12β is a predominant isoform in adult tissues^40^ and a pivotal modulator of pathologies ranging from cancer to neurodegenerative disorders and autoimmune diseases.^41–44^ Upon binding its receptors, CXCR4/CXCR7, CXCL12β drives tumor progression by promoting angiogenesis, epithelial-to-mesenchymal transition, and organ-specific metastasis.^44–47^ Furthermore, this axis induces resistance to chemo- and radiotherapy^48,49^ and can blunt immunotherapy efficacy by modulating immune cell populations within the tumor.^50,51^ By silencing CXCL12β, LIV may cripple this CXCL12β-CXCR4/CXCR7 signaling axis, depriving cancer cells of pro-survival and pro-motility cues that underpin their systemic dissemination. This positions LIV as a novel mechanobiological intervention for disrupting protumorigenic communication between cancer cells and their microenvironment, and inhibiting the metastatic potential of aggressive breast cancers.^52,53^

Our data indicates that LIV induces a genome-wide suppression of MDA-MB-231 transcriptomes, a pattern visually confirmed by a comprehensive heatmap of differentially expressed genes (Figures 5E, 5F and 5G). This broad transcriptional repression appears to be orchestrated by a coordinated shift in miRNA expression. Specifically, LIV induced the upregulation of two anti-tumor miRNAs: hsa-miR-141, which suppresses growth and metastasis by targeting MXRA8,^54^ and hsa-miR-122, a known tumor suppressor that blocks proliferation and tumorigenesis by targeting IGF1R.^55^ Concurrently, LIV downregulated protumorigenic miRNAs, including hsa-miR-105-5p, a promoter of tumor growth and metastasis,^56–58^ and hsa-miR-181a-5p, an oncogene that inhibits SPRY4 and correlates with poor patient outcomes,^59,60^ Collectively, these findings demonstrate that LIV enforces a coordinated, miRNA-mediated transcriptional brake that disrupts oncogenic and pro-inflammatory gene networks. GO enrichment showed that DEGs were predominantly localized to nuclear and cytoplasmic compartments, such as nucleoplasm, chromatin, and cytoskeleton. In parallel, KEGG pathway analysis identified significant enrichment in oncomiRs, pan-cancer signaling, cytoskeletal control, breast cancer-specific pathways, and mechanotransduction cascades.

Mirroring the scheduled dosing regimens characteristic of pharmacological interventions, LIV-induced effects may too, be ephemeral, necessitating regular redosing. But unlike pharmacological interventions—which are often limited by cumulative toxicity and metabolic byproducts—LIV leaves no chemical residue, allows for tissue recovery between sessions, and can be safely administered without dose-limiting side effects.

In summary, our findings demonstrate that LIV either fully or partially suppresses key inflammatory mediators, IFNγ, CXCL-12β, and IL-6, while promoting or sparing resolving mediators, MCP-1 and IL-10. This reflects a broader, mechanically-induced transcriptomic shift that wires mechanosensitive signaling hubs to disease-relevant networks controlling cell fate and intercellular communication. Therapeutically, LIV orchestrates gene regulatory networks and miRNA expression to suppress oncogenic signaling, resolve inflammation, and curb malignancy.

## RESOURCE AVAILABILITY

### Lead contact

Requests for additional information and resources should be addressed to and will be provided by the lead contact, Ray P.S. Han

### Materials availability

This research study did not produce any new unique reagents.

### Data and code availability

The bulk RNA sequencing datasets created for this research are archived in GEO Submission (GSE316980), which are accessible to the public as of the publication date. All other experimental data mentioned in this paper will be provided by the lead contact upon request, after confirming any intellectual property or confidentiality obligations. The mechanical stimulation of cultured cells and biomolecular analyses were conducted as outlined in STAR Methods. Additional details will be provided by the lead contact upon request.

## ACKNOWLEDGMENTS

This work was supported by Jiangxi University of Chinese Medicine Innovation Team Funding (Grant No. CXTD22018) and Jiangxi University of Chinese Medicine Cancer Research Centre Start-up Funds (Grant No. 12418008).

## AUTHOR CONTRIBUTIONS

This work was a collaboration among researchers from two Nanchang universities, The Ocean Road Cancer Institute of Tanzania, and medical practitioners from three Nanchang hospitals. We express our gratitude to all team members for their support throughout this project. The specific contributions are as follows: conception, design, and methodology development—S.S.M., R.H.; data acquisition— S.S.M., B.K.K., G.F.S., X.H., C.H., W.Z., Y.Z., S.Y.; imaging, data analysis, and interpretation—S.S.M., B.K.K., G.F.S., B.L.; administrative, technical, and material support—S.S.M., L.W., J.M., X.H., R.H.; writing and manuscript revision—all authors; funding acquisition and supervision—R.H.

## STAR★METHODS

Detailed methods are provided in the online version of this paper and include the following:

- KEY RESOURCES TABLE
- EXPERIMENTAL MODEL

- Ethics statement
- Cell line
- Cell culture models
- METHOD DETAILS

- Design and calibration of a mechanical vibrator system
- Creation of tumor-immune microenvironment (TIME)
- Spheroid formation and assay
- Cell viability assay
- Cell migration assay
- Scratch wound gap area analysis by ImageJ
- Extracellular vesicle extraction
- Nanoparticle tracking analysis (NTA)
- Zeta potential analysis
- Confocal Laser Scanning Microscope (CLSM) imaging
- Transmission electron microscope
- ELISA assay
- QUANTIFICATION AND STATISTICAL ANALYSIS

- RNA sequence analysis
- Genomic DNA isolation and qRT-PCR analysis
- Other statistical analyses

## SUPPLEMENTAL INFORMATION

Supplemental information can be found online at https://doi.org/………………

Received:

Revised:

Accepted:

## STAR★METHODS

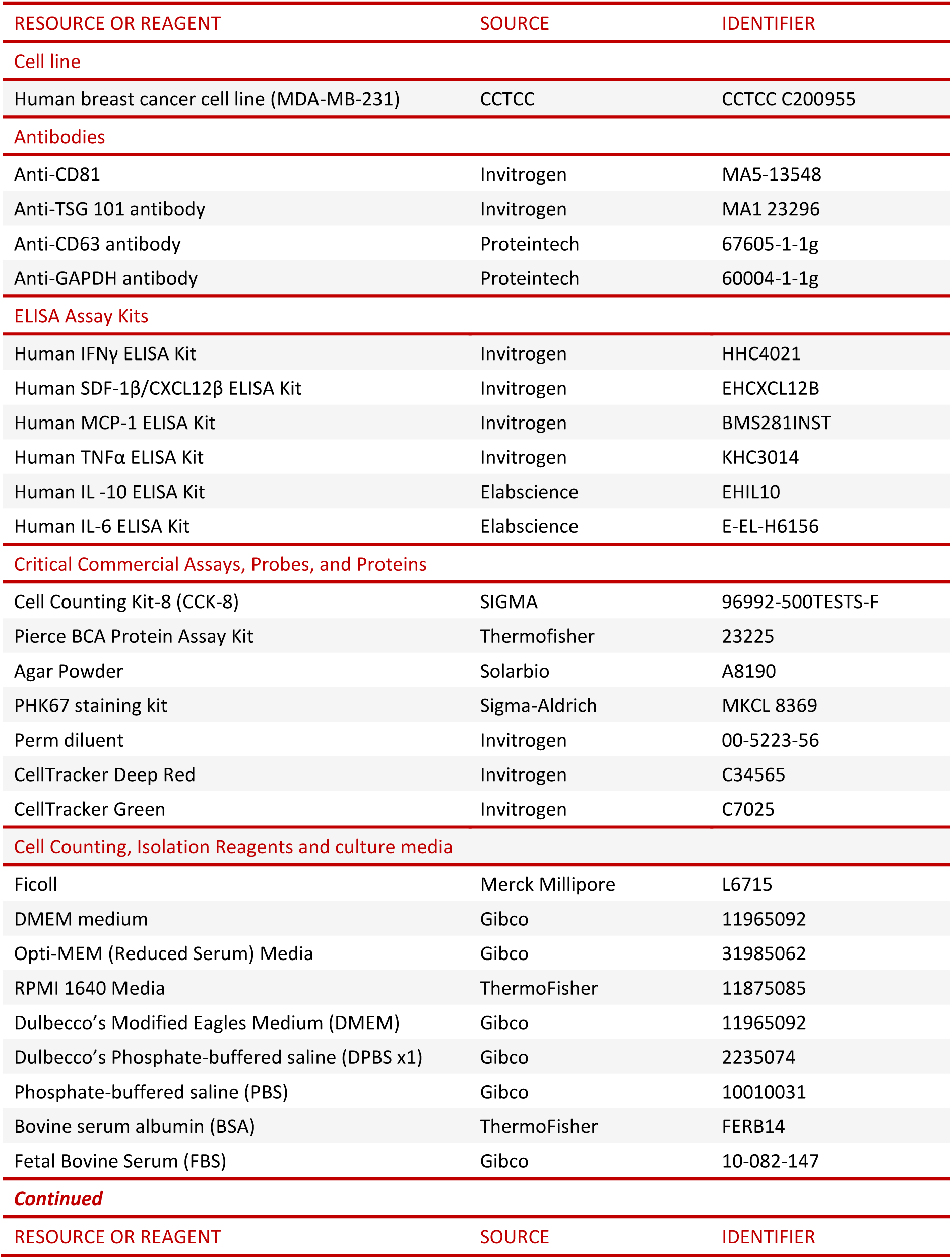
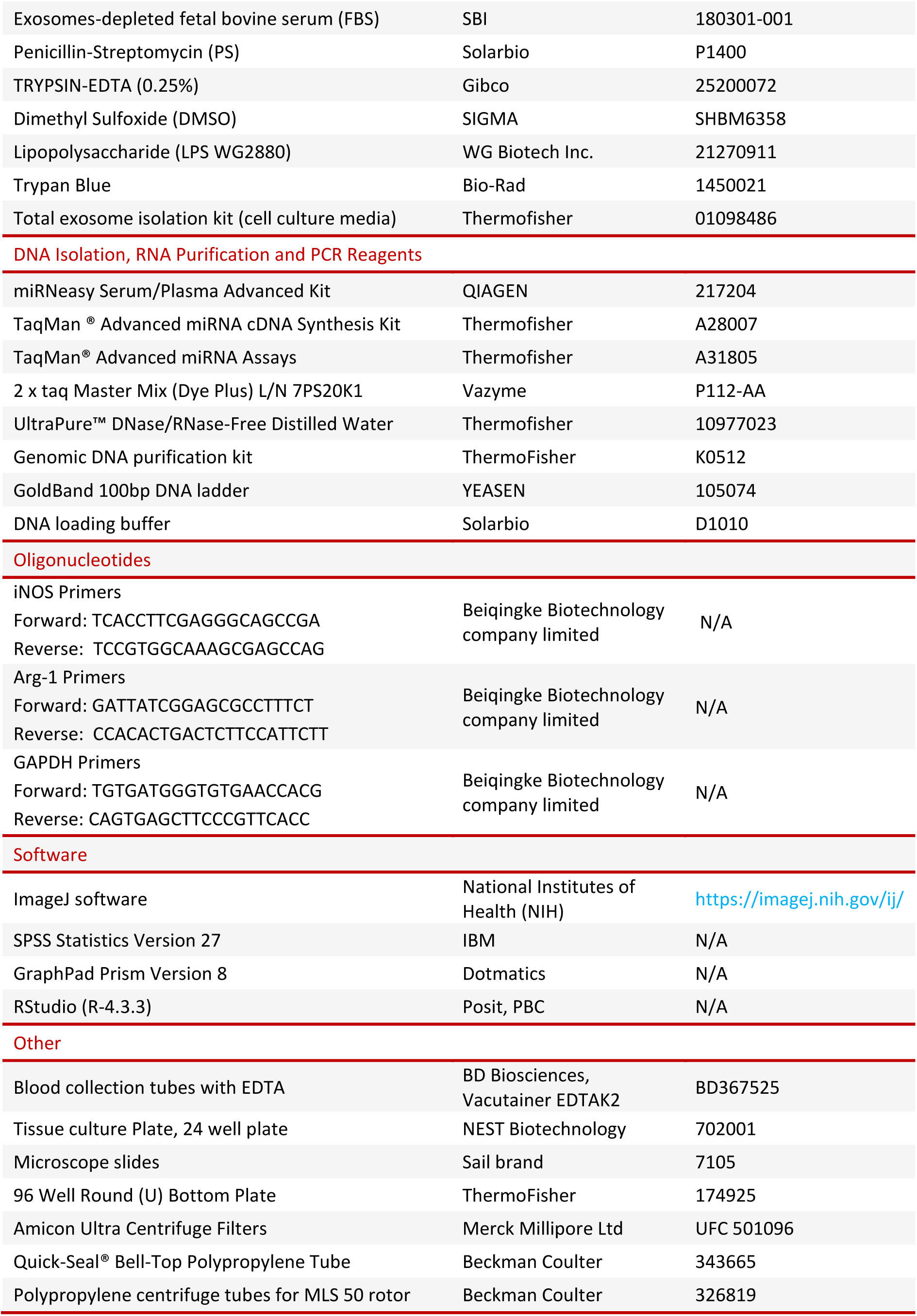
KEY RESOURCES TABLE

## EXPERIMENTAL MODEL AND STUDY DESIGN

### Ethics Statement

This study adheres to all applicable ethical standards. Written informed consent from participants was obtained after Institutional Review Board approval (IRB no. JZFYLL202202230010) for collecting whole blood samples from five healthy volunteers for the purpose of isolating PBMCs.

### Cell line and culture model

The human breast cancer cell line MDA-MB-231 was purchased from the China Center for Type Culture Collection (CCTCC), in Wuhan. The MDA-MB-231 cells were maintained in DMEM medium (Gibco™ 11965092), supplemented with 10% FBS (10-082-147) and 1% Penicillin-Streptomycin (P1400). Human PBMCs were isolated from heparinized whole blood by using Ficcol Paque PLUS (L6715) solution-based density gradient centrifugation as per established protocol.^61^ PBMCs were cultured at a density of 5 x 10^5^ cells per mill in RPMI-1640 medium (11875085) and their activation was stimulated with LPS (WG2880). Both MDA-MB-231 cells and PBMCs were incubated in a humidified incubator with 5% CO_2_ at 37°C. In addition, we created a 3D co-culture platform that bio-mimic in vivo tumor-immune microenvironment by co-culturing MDA-MB 231 cells and PBMCs. The protocol described by Saraiva et al.^61^ was adopted in generating spheroids (3D co-culture) that are biomimetic of the tumor-immune microenvironment.

### Design and calibration of a mechanical vibrator system

A mechanical vibrator system was developed to investigate the impact of mechanical stimulation within the tumor-immune microenvironment (TIME) on the management of CPI and metastasis. The vibration system is composed of two components: a vibration device and a rigid vibration support system (stand) constructed from steel bars, as illustrated in Figures 9A, 9B. The vibration device features six settings that can generate a specific vibration signal strength with frequencies that vary from 90 Hz to 300 Hz for the cultured cells. The stand offers both stability and flexibility to the vibrator, enabling it to deliver consistent/reproducible mechanical vibration cues.

A portable digital vibrometer (Figure S9C), equipped with a piezoelectric transducer, was utilized to calibrate the system, ensuring that the cultured cells are exposed to the appropriate mechanical signal that corresponds to the desired vibration frequencies. The vibrometer employed in our study is a smart sensor (version number 6-AS63A-1216-02), which utilizes a piezoelectric acceleration transducer to transform the mechanical vibration signal into an electrical signal. This vibrometer can measure an acceleration frequency range of 10 Hz to 15 kHz.

During the calibration process, an equal volume of culture media was introduced into the tissue culture dish to replicate the actual cell culture environment. The culture plate was secured at the base of the vibration system using a tissue plate stabilizer to minimize movement during vibration. As a result, the vibrometer probe was positioned at the bottom of the tissue plate, and the vibrator was activated, followed by the recording of vibrometer measurements. Various vibration signal strengths corresponding to different gears provided by the vibrator were documented from the vibrometer readings. For each gear, a minimum of three readings were collected, and averages were calculated to ascertain the exact vibration signal strength in terms of frequencies (Hz).

Cultured PBMCs were stimulated with 5 µg/ml LPS to induce an inflammatory response. Following this, the CPI was created by co-culturing breast cancer cells (MDA-MB-231) with LPS-stimulated PBMCs exhibiting a pro-inflammatory phenotype. Importantly, both MDA-MB-231 cells and PBMCs were grown in either monoculture or co-culture systems using exosome-depleted media. To achieve a suitable mechanical signal strength that could suppress the secretion of PICs, the cultured cells were subjected to vibrations at frequencies of 90 Hz, 150 Hz, and 300 Hz for 15 minutes over three consecutive days. The PICs were measured from the collected cell culture supernatant, and the vibration frequency that resulted in the lowest concentration of PICs was identified. To ascertain the optimal LPS concentration for stimulating PBMCs, cultured PBMCs were treated with varying concentrations of LPS (0, 5, 20 & 40 µg/ml), and an evaluation of cell responses was conducted by measuring viability and quantifying levels of IL-6 cytokine, a PIC.

Furthermore, the appropriate timing for collecting the cell culture supernatant to measure PICs was determined by stimulating PBMCs with 5 µg/ml LPS and assessing the PIC concentration at 12, 24, and 48 hours. The cell culture supernatants derived from vibrated cells (experiment) and non-vibrated cells (control) were stored at -200 °C for extended durations or at -80 °C for shorter durations. We examined cell migration as a metric for evaluating cancer spread/metastasis. To investigate the influence of LIV on cell migration, we employed a scratch wound healing assay technique to ascertain if the mechanical signal could either diminish or promote cell migration. Consequently, we co-cultured MDA-MB-231 cells with LPS-stimulated PBMCs (Figure S10) to establish a TIME for evaluating the impact of LIV on cell migration. The cells were categorized into two groups: the experimental group (vibrated cells) and the control group (non-vibrated cells). Scratch wounds were generated using a 200 µL pipette tip (Figure S2) in accordance with the protocol outlined by Justus et al.^62^ Images were captured at baseline (0 hours), followed by assessments at 6 hours and 12 hours to evaluate the closure of the wound gap (Figure S3). The potential for migration was quantified by measuring the area of the scratch wound gap with ImageJ software, and comparisons were made between the experimental group and the control group (Figure S4).

We performed a co-culture of MDA-MB-231 cells and PBMCs in Transwell to evaluate the impact of mechanical stimulation on cellular responses through gene expression and subsequent protein synthesis. This co-culture technique enables cells to communicate via the transfer of signaling biomolecules, despite their physical separation, and permitted the individual characterization of cells as well as the extraction of genomic DNA for further molecular analysis.

As a result, the 2D co-culture experiments were conducted using 6-well Transwell plates. These plates are fitted with permeable Transwell inserts that create two separate platforms for cell cultures. The base of the 24mm Transwell insert is designed to be permeable, featuring a polyester membrane with pore sizes of 0.4 μm, which promotes the exchange of signaling biomolecules and EVs between cells in the distinct co-culture platforms.

The MDA-MB-231 cells were cultured at a density of 1 x 10^6^ cells/ml in the upper platform of a Transwell insert one day prior to culturing PBMCs at a density of 4 x 10^6^ cells/ml on the lower platform of a Transwell. Initially, MDA-MB-231 cells were maintained in DMEM complete medium, but the culture medium was changed to RPMI complete medium the following day to support the growth of PBMCs. Two Transwell plates were utilized for co-cultivation following a similar protocol, with one plate designated for the vibration experiment and the other for the control. After three days of culture, genomic DNA was extracted from the PBMCs for gene expression profiling. We also evaluated macrophage polarization by measuring iNOS and Arg-1 gene expression, which are indicative of M1 and M2 phenotypes, respectively, while GAPDH served as the house control gene. The primer sequences listed in Extended Data Table 3 were employed to assess the expression levels of the iNOS, Arg-1, and GAPDH genes, which were procured from Beiqingke Biotechnology Company Limited, Hangzhou branch.

### Cell viability assay

MDA-MB 231 cells were cultured in two 6-well plates, one designated for the experimental group and the other for the control group. Once the cells achieved 70-80% confluence, they were mechanically stimulated with a vibration frequency of 90 Hz for a duration of 15 minutes over three consecutive days. Following this, the culture medium from both the experimental and control dishes was discarded, and the cells were rinsed twice with warm PBS (1x) prior to being detached using trypsin-EDTA (0.25%). The cells were then counted with a hemocytometer and subsequently plated into the corresponding 96-well plates for both the experimental and control conditions. In both groups, 1,000 cells were suspended in DMEM culture media at a volume of 100 μl per well and pre-incubated at 37°C with 5% CO_2_ for 12 hours (overnight). The following morning, 10 μl of Cell Counting Kit-8 (SIGMA, item number: 96992-500TESTS-F) solution was carefully added to each well of the 96-well plate without creating bubbles, and the plates were incubated for 4 hours at 37°C with 5% CO_2_. Subsequently, the absorbance at 450 nm of the 96-well culture plates was assessed using a microplate reader. The optical density (OD) values obtained were utilized for the statistical analysis of the gathered dataset. The calculation of the percentage of viable cells was performed using the following formula: Cell viability (%) = [(As-Ab)/(Ac-Ab)] x 100. In this equation, As represents the absorbance of the experimental well, which includes the absorbance of vibrated cells, culture medium, and CCK8. Conversely, Ac denotes the absorbance of the control well, which consists of absorbance from wells containing non-vibrated cells, culture medium, and CCK8, while Ab signifies the blank absorbance, which is the absorbance from wells that contain only culture medium and CCK8.

### Extracellular vesicles extraction

The total exosome isolation kit (01098486) was utilized to extract exosomes from cells that were cultured in media depleted of exosomes, following the protocol provided by the manufacturer. The reagent was incorporated into the supernatant at a volume ratio of 1:2, and the mixture was vortexed to achieve a uniform solution. The supernatant was carefully aspirated without disturbing the exosome pellets and was subsequently discarded. Following this, the exosome pellets were suspended in 200 µl of PBS x1.

### Nanotrack Analysis (NTA)

The NTA of EVs was conducted according to the method established by *Martin E.M. Parsons* and colleagues.

### Zeta potential analysis

The NanoBrook Omni-Particle Zeta Potential Analyzer was employed to assess the surface charge of EVs derived from samples. The electrode plate and cuvettes underwent two rinses with deionized water and were subsequently dried using a non-abrasive tissue. The EV sample was suspended in a low-ionic-strength medium devoid of Ca^2+^ and Mg^2+^. Following this, the zeta potential was measured and analyzed utilizing Particle Solutions v. 3.6.0.7122 software. The parameters for the sample were calibrated to a pH of 7, a viscosity of 0.890, and a refractive index of 1.331. The selected model for zeta potential was based on Smoluchowski equation.^63^

### Confocal laser scanning microscopic (CLSM) imaging

CellTracker red (C34565) and green (C7025) were used to stain cells, while EVs were stained with PKH67 green fluorescent dye (MKCL 8369). The lower magnification objective lenses of 20x and 40x were utilized for cell imaging, whereas a higher magnification objective lens of 60x from a Leica Lesser Scanning Confocal Microscope (LSCM), along with the use of immersion oil, was selected for the acquisition of images of EVs. The excitation wavelengths for far-red CellTracker, green CellTracker, and green PKH67 fluorescence were 637 nm, 521 nm, and 488 nm, respectively. The final images were processed using the Imaris software.

### Transmission electron microscope (TEM) imaging

The EV sample intended for TEM evaluation was prepared following the guidelines provided by Théry et al.^64^

### Western blotting

MDA-MB-231-derived EVs were lysed using RIPA buffer (SKU: 89900) and quantified with the Pierce Protein Assay BCA kit (catalogue number 23225) following the manufacturer’s instructions. The extracted protein sample was subjected to heating at 95 °C, subsequently loaded into a running gel for protein separation, and transferred to a PVDF membrane. The PVDF membrane was then blocked with a 5% skim milk solution.

Primary antibodies targeting the tetraspanin proteins of EVs were evaluated using anti-CD63 (67605-1-1g), anti-CD81 (MA5-13548), anti-TSG 101 (MA1 23296), and anti-GAPDH antibody (60004-1-1g). As a result, the primary antibodies were incubated with the PVDF membrane overnight. Finally, the PVDF membrane was washed five times and incubated with a secondary antibody (HRP-conjugated, SA00001-1) for two hours at room temperature. Following the incubation, any unbound secondary antibody was eliminated by washing six times with TBST buffer (x1). The PVDF membrane, which contained the target protein band, was bound to the HRP secondary antibody, which produced a fluorescent signal upon the addition of HRP substrate (34075, ThermoFisher). The fluorescent signal was then visualized using a chemiluminescent immunoassay analyzer.

### ELISA assay

The sample for both the experimental and control groups was obtained from serum-free cell culture supernatant and was prepared in accordance with the instructions provided by the manufacturer of the commercial human ELISA kits. The kits utilized included the human IL-6 ELISA Kit from Elabscience (E-EL-H6156), the IFN-γ ELISA kit from Invitrogen (HHC4021), the CXCL12β ELISA Kit from Invitrogen (EHCXCL12B), the MCP-1 ELISA Kit (BMS281INST), the TNFα ELISA Kit from Invitrogen (KHC3014), and the IL-10 ELISA Kit from Elabscience (EHIL10).

### Genomic DNA isolation and PCR analysis

Genomic DNA (gDNA) was isolated following the guidelines provided by the manufacturer of the ThermoFisher genomic DNA purification kit (reference number: K0512). The specific sequences of iNOS, Arg-1, and GAPDH (10 μM) were amplified from the extracted gDNA (> 1ng). The PCR protocol used to assess the expression levels of the genes of interest adhered to the methodology outlined by Lorenz^65^ and agarose gel electrophoresis was performed as previously described by Lee et al.^66^

### Micro-RNA Isolation and real time-PCR (qRT-PCR)

The isolation of micro-RNAs from EVs derived from MDA-MB-231 was accomplished utilizing the miRNeasy Serum/Plasma Advanced Kit from Qiagen (catalogue number 217204) in accordance with the manufacturer’s instructions. Subsequently, cDNA was generated from miRNA employing the TaqMan Advanced miRNA cDNA Synthesis Kit (A28007). The detection and quantification of PCR were carried out using TaqMan Advanced miRNA Assays (A31805) and TaqMan Fast Advanced Mastermix (4444556), which are specifically chosen primer and probe sets intended for the analysis of expression levels of particular micro-RNAs (miRNAs) of interest. Further, BGI Genomics conducted sequencing of samples on the DNBSEQ platform (https://en.genomics.cn/), whereby samples were sequenced in triplicate on the DNBSEQ platform and Agilent Technologies were used for RNA library preparation. The differential expression analysis of miRNAs was determined if DEseq2 for [log2FC]> 0 & Qvalue< 0.05, or if DEGseq and PossionDis [log2FC]= 1 & Qvalue< 0.001.

### Statistical Analysis

Statistical analyses were performed using IBM SPSS Statistics Version 27, GraphPad Prism Version 8 and RStudio (R-4.3.3). Data are expressed as the mean± standard deviation (S.D.) of three or more individual experiments conducted in triplicate. The comparison of means between two data sets was assessed using a 2-tailed Student’s unpaired t-test at a 95% confidence level. A significance level was established when the *p*-value is below 0.05.

## Supplemental figures and tables

**Figure S1.**
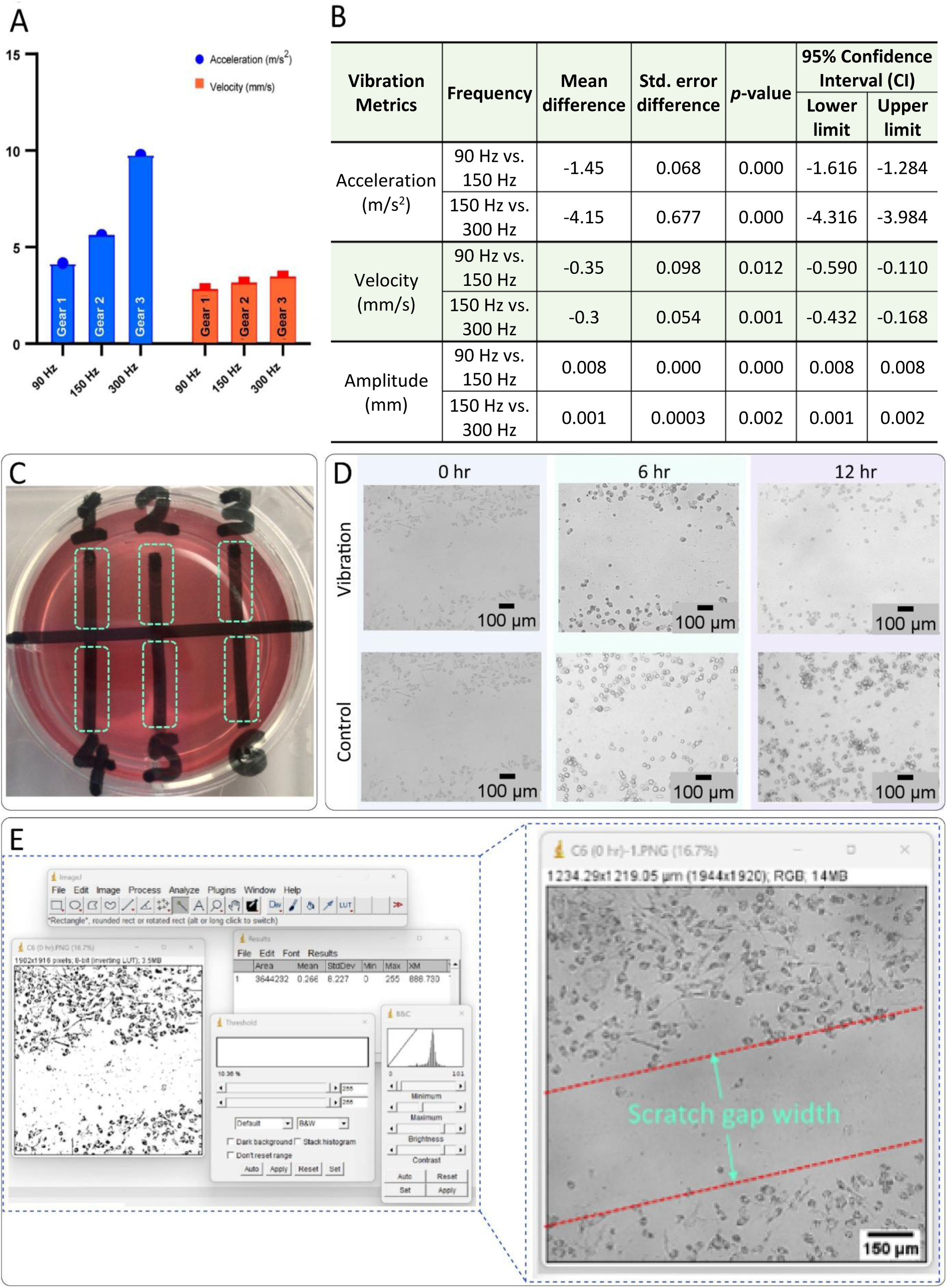
Calibration of the vibration system and the wound scratch analysis. (A) Calibration results showed that an increase in the number of gears of the vibration motor correlated with an increase in the mean vibration acceleration and velocity (*n* = 4, *p* < 0.04). Related to Figure 1C. (B) Statistics for multiple calibration metrics: (a) acceleration, (b) velocity, (c) amplitude, related to Figure 1C. (C) A cell culture well (with culture media) labeled with black vertical lines 1 to 6 representing scratch wounds (dotted green lines) and the reference horizontal line, which helped to reposition the plate in the microscope. Related to Figure 2F-H. (D) Scratch wound images of co-cultured MD MB-231 cells and PBMCs taken at the start 0 h, 6 h, and 12 h. The same image at 0 h was used as a reference image to compare with wound closure in both vibration and control at different time points. Related to Figure 2F-H. (E) Scratch wound gap area analysis by ImageJ, (https://imagej.nih.gov/ij/), related to Figure 2F-H.

**Figure S2.**
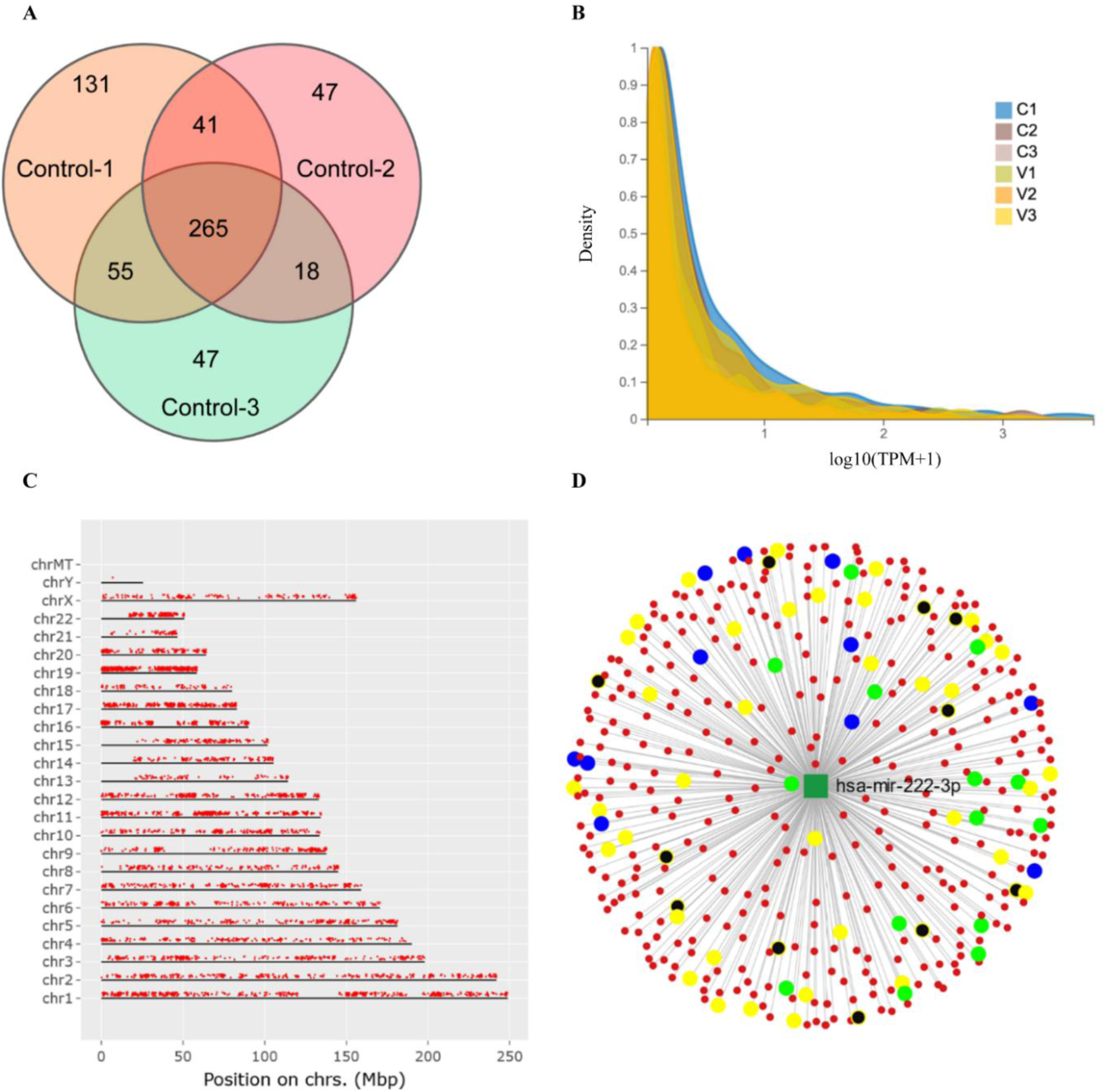
Correlation of gene sets expressed in control samples, global density map of RNA, gene mapping, and number of transcripts regulated by individual microRNA. Related to Figure 5 (A) VENN diagram depicting the correlation of the gene set expressed in control samples (C1, C2, C3). Each circle represents a group of gene sets of individual samples, and the areas overlaid by different circles represent the intersection of gene sets among samples and the non-overlapping part indicates genes that are uniquely expressed. (B) Density map of RNA representing each sample, where the prefixes C and V in the legend denote control and vibration, respectively. (C) Gene map depicting the location of our target genes on chromosomes (chrs). (D) miRNet network analysis of GO terms in the biological processes (BP) category regulated by hsa-miR-222-3p. The nodules represent transcript, mostly in red with specific groups of transcripts represented by different colors. Yellow indicates transcripts associated with cytoskeletal regulation, green represents transcripts associated with apoptotic signaling, blue depicts transcripts related to translation regulation and black shows transcripts related to regulation of other genes.

**Figure S3.**
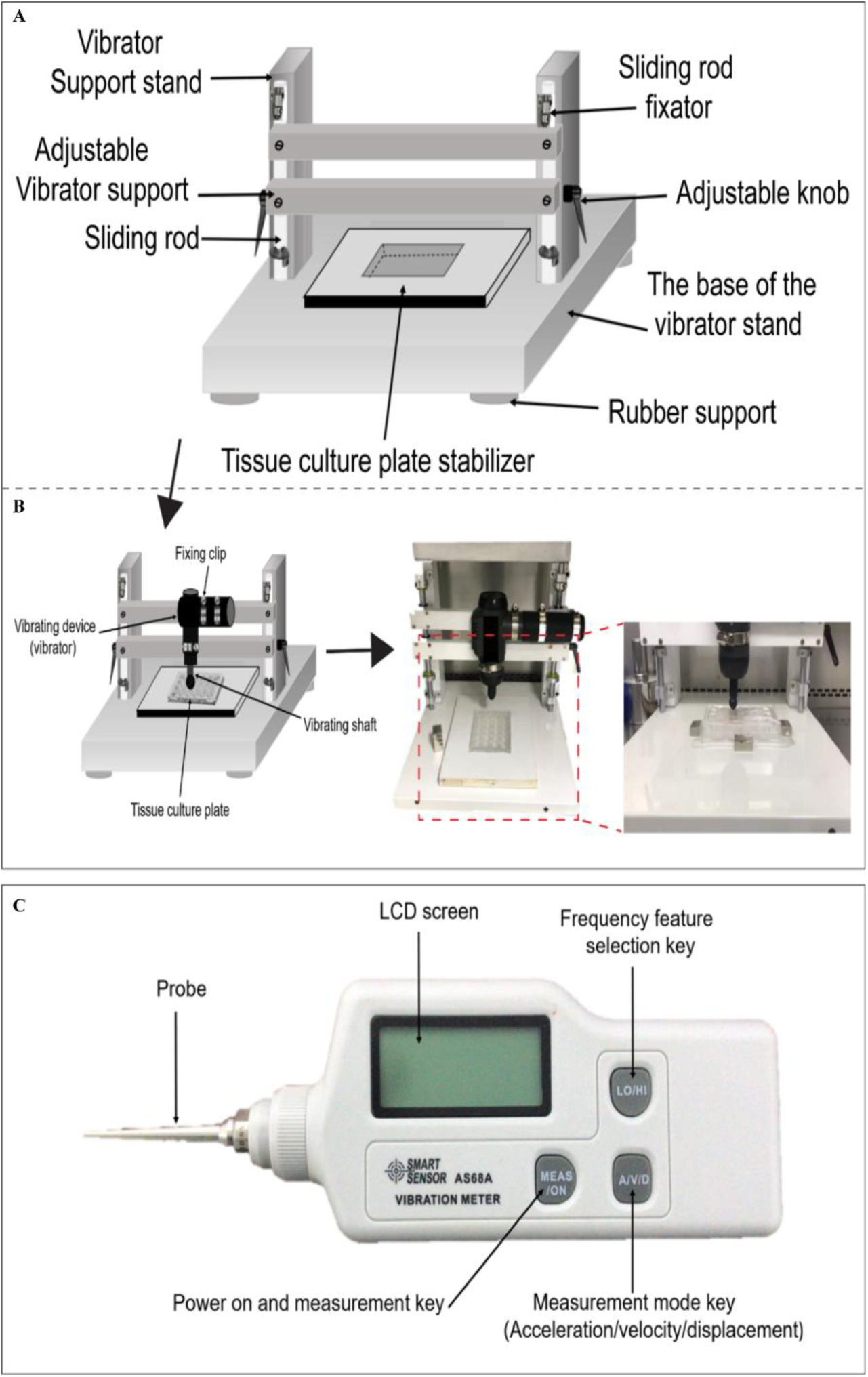
Layout of the vibration system and vibrometer, related to. Figure 1A (A) Rigid vibrator stand. (B) Massage gun (vibrator) fixed in a stand for loading-controlled mechanical vibration cues. (C) Vibrometer used to calibrate mechanical signal.

**Figure S4.**
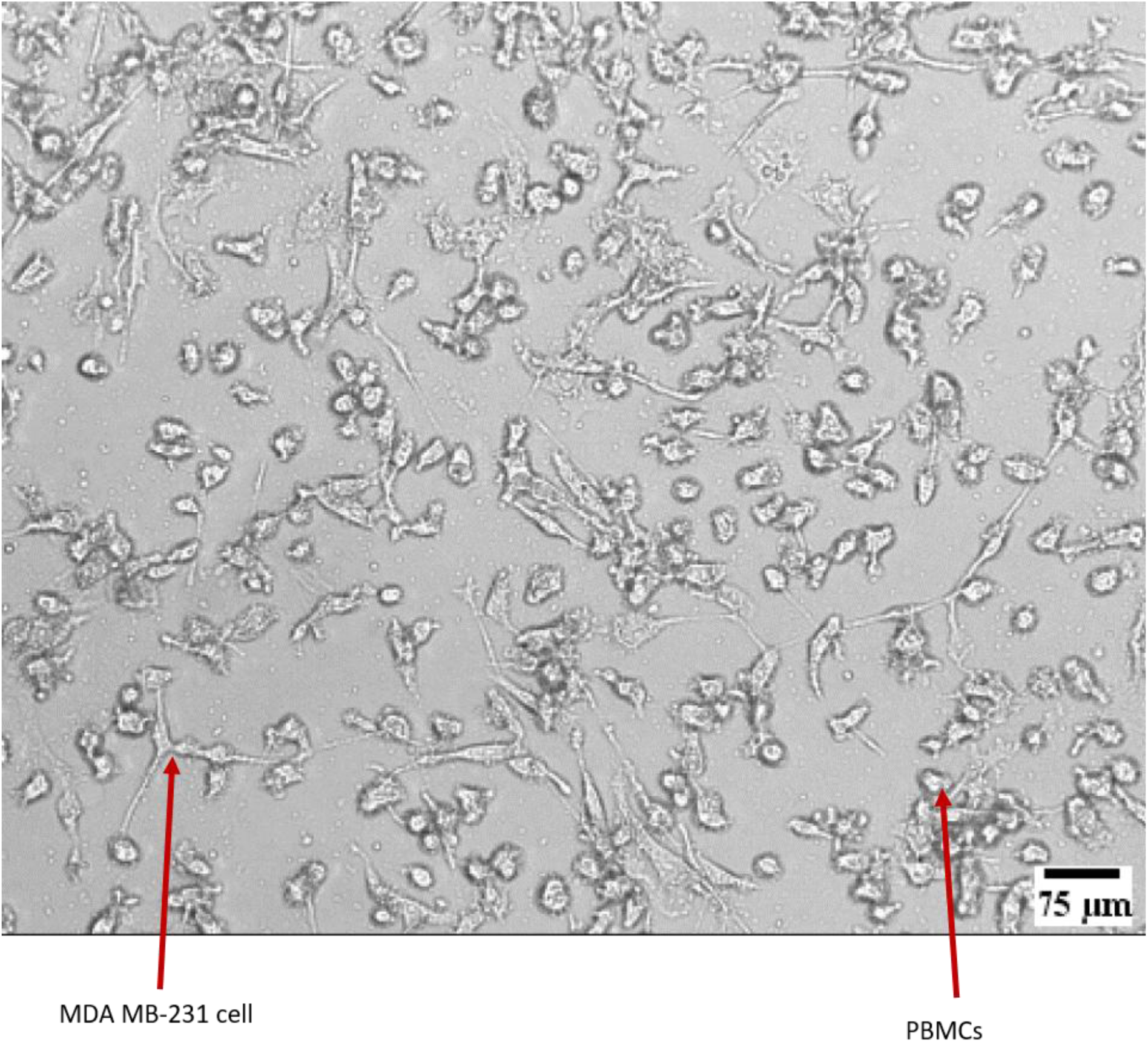
Monolayer co-culture of MDA-MB-231 cells and PBMCs, related to Figure 2E

**Table S1.**
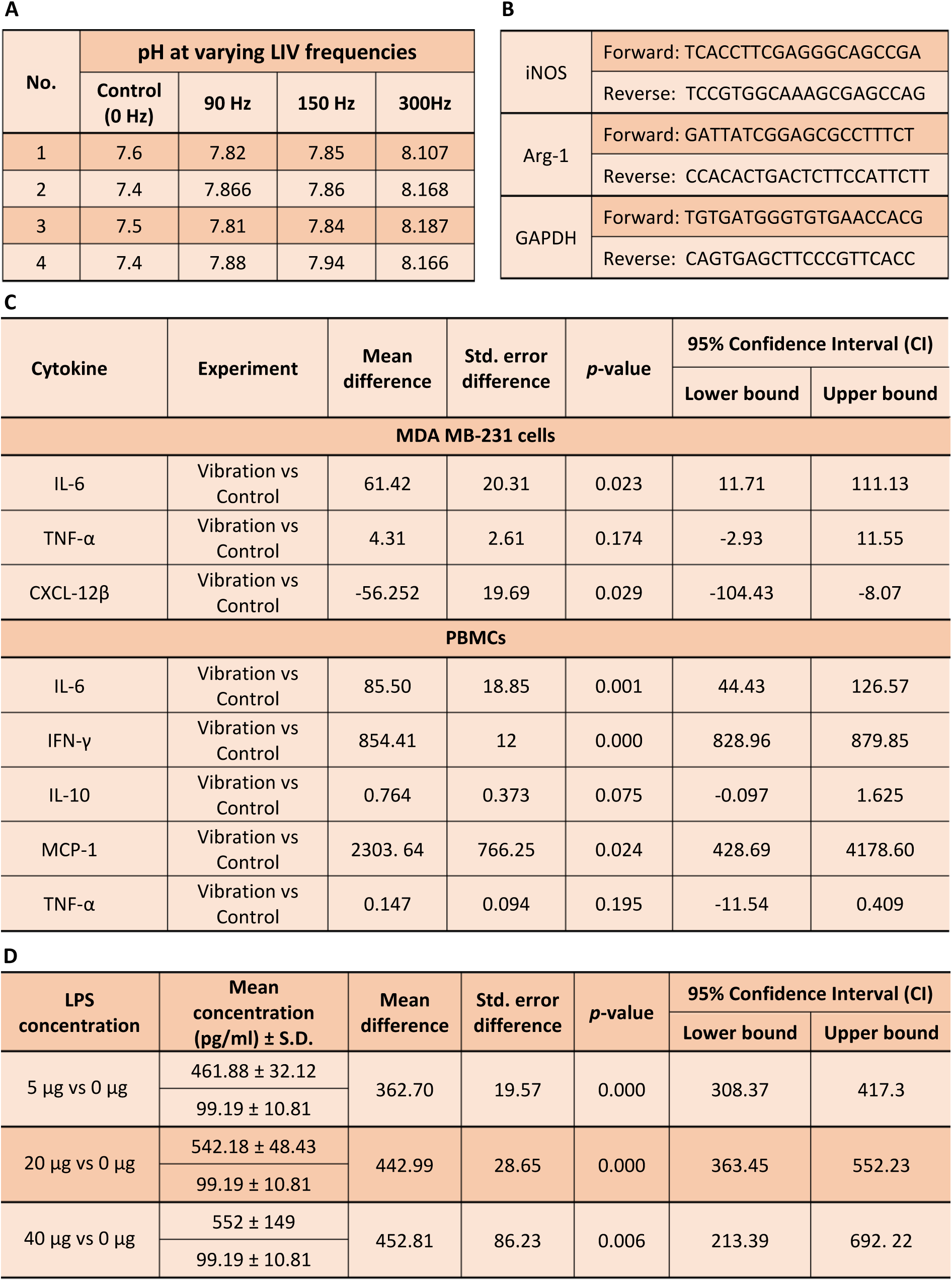
pH readings of supernatant from cell cultures, primer sequences for macrophage genes, and consolidated statistics from ELISA assay (A) pH levels measured in the supernatant of LPS-stimulated PBMCs following exposure to different LIV frequencies (*n* = 4), related to Figure 1G. (B) Primer sequences of iNOS, Arg-1, and GAPDH genes, related to Figure 2M-P. (C) ELISA assay statistics for comparisons between vibration and control groups, related to Figures 3I, J. (D) Statistics of IL-6 assay for PBMCs stimulated with different concentrations of LPS, related to Figures 3I, J.

**Table S2.**
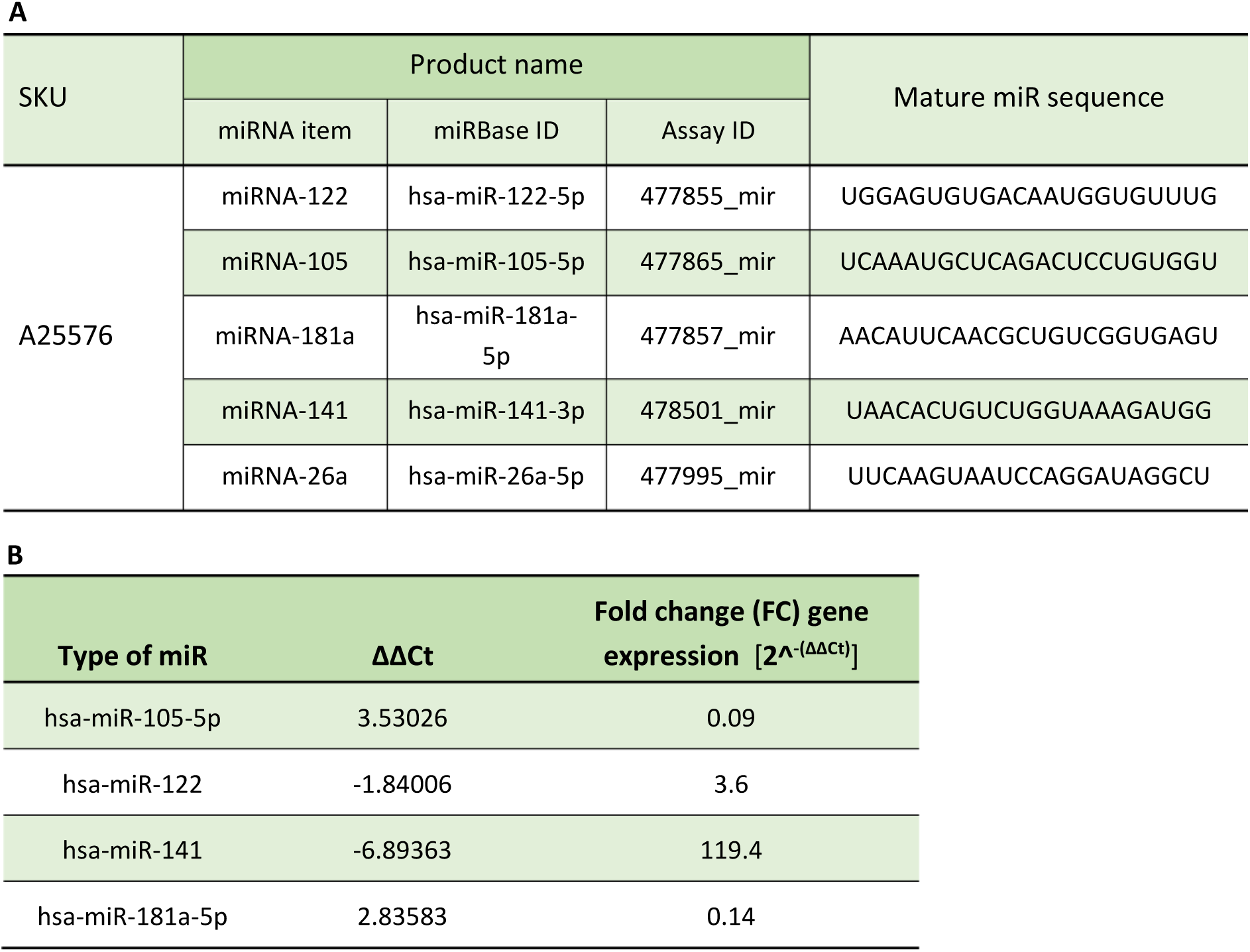
Primer sequences for specific genes and varying expression levels of targeted microRNA. (A) TaqMan primer sequences for various miRs, related to Figure 5H. (B) Differential expression levels of miR expressed as relative fold-change (FC) gene expression (control vs. experiment), related to Figure 5H.

